# Shared ligand-blocking mechanism but distinct conformational modulation by α5-targeting antibodies BIIG2 and MINT1526A

**DOI:** 10.1101/2025.01.08.631572

**Authors:** Adam Nguyen, Joel B. Heim, Gabriele Cordara, Matthew C. Chan, Hedda Johannesen, Cristine Charlesworth, Ming Li, Caleigh M. Azumaya, Benjamin Madden, Ute Krengel, Alexander Meves, Melody G. Campbell

## Abstract

Integrins are heterodimeric receptors important for cell adhesion and signaling. Integrin α5β1 is a key mediator of angiogenesis and its dysregulation is associated with tumor progression and metastasis. Despite numerous efforts, α5β1-targeting therapeutics have been unsuccessful due to poor efficacy and off-target effects. A contributing factor is our limited understanding of how integrin conformation influences interactions with therapeutics. Using cell-based functional assays, patient-derived xenografts, biophysics, and electron microscopy, we shed light on these relationships by characterizing two anti-α5β1 antibodies, BIIG2 and MINT1526A. We show that both antibodies bind α5β1 with nanomolar affinity, reduce tube formation *in vitro*, and bind overlapping epitopes that block fibronectin binding. However, using electron microscopy, we reveal that while BIIG2 binding does not substantially alter the conformational states, MINT1526A preferentially recognizes the bent conformation and restricts the conformational ensemble. These insights can guide which aspects to prioritize to improve the design of future integrin-targeted therapeutics.

## Introduction

Integrin heterodimers, which consist of an α subunit and a β subunit, are type I transmembrane receptors that play vital roles in immune response, cell motility, cell-matrix interactions, signaling, proliferation, and apoptosis ^1–3^ (**Figure S1A**). Pathologies linked to integrins include autoimmune, cardiac, pulmonary, neurodegenerative, hematologic, and oncologic conditions ^4,5^. Integrins have long been identified as promising therapeutic targets. Although numerous integrin-targeting therapeutics have entered clinical trials, fewer than a dozen have been approved and marketed, while others have been withdrawn due to limited efficacy and adverse effects ^6,7^.

Integrin α5β1 plays a key role in facilitating cell migration during angiogenesis and is one of several integrins that recognize the conserved arginine-glycine-aspartate (RGD) sequence motif found in several extracellular matrix (ECM) proteins. The primary α5β1 ligand, fibronectin (FN), is a glycoprotein that scaffolds other matrix proteins and cells. α5β1 participates in tumor angiogenesis, among other pathologically significant processes related to human diseases ^8–10^. Like many other well-studied integrins, α5β1 adopts three conformations which can be directly linked to α5β1 activation: bent, extended closed, and extended open (**Figure S1B**). The extended open conformation is stabilized by FN via two interactions: the canonical integrin binding RGD motif and an additional synergy site. Specifically, a fibronectin domain (type III domain 10, FN10) contains an RGD motif that inserts into the α and β subunit interface ^11,12^ and a second domain, FN9 contains the synergy site that facilitates a catch-bond interaction with the β-propeller of the α5 subunit ^12–16^.

The increased expression of integrins α5β1, αvβ3, and αvβ5 in new blood vessels during the angiogenic switch in tumors underscores their significance as potential targets for therapeutic intervention ^4,17–19^. Early-phase clinical trials involving patients with advanced solid tumors investigated multiple monoclonal antibodies targeting α5β1 function, however, these trials did not demonstrate clear efficacy ^20–22^. Among the tested antibodies was MINT1526A (RG-7594), a fully humanized IgG1 antibody specifically formulated to block α5β1 function ^22^. A different α5β1-inhibiting antibody, BIIG2, was developed to inhibit human JAR choriocarcinoma cells binding to ECM proteins by immunization of Lewis rats ^23^. Follow-up studies showed that BIIG2 was effective in both inhibiting fibronectin attachment and identifying invasive cytotrophoblasts within the human placenta ^23,24^. To better understand how integrin conformation influences antibody-mediated inhibition, we sought to characterize BIIG2’s functional and mechanistic properties. We compare it to MINT1526A to highlight differences that could contribute to antibody effectiveness, since MINT1526A was well tolerated in patients during phase I clinical trials but was shown to have limited advantages over existing therapies ^22^. Therefore, we evaluated BIIG2 and MINT1526A using structural, biophysical, and cell-based methods to define the mechanisms underlying antibody-mediated α5β1 inhibition. We found that BIIG2 shows more favorable inhibitory outcomes *in vitro* when compared to MINT1526A, and the two antibodies elicit different effects on conformations accessible to α5β1. Since integrin conformation has been shown to influence the severity of off-target effects ^6,7^, our results provide the molecular basis for the design of future integrin-targeted therapeutics.

## Results

### BIIG2 and MINT1526A antibodies reduce endothelial tube formation in vitro

Angiogenesis is the formation of new blood vessels and has been described extensively as critical for tumor growth ^8,25^. To evaluate whether BIIG2 or MINT1526A can inhibit angiogenesis in an integrin α5β1-dependent manner, we employed a two-dimensional tube formation assay by cultivating human umbilical vein endothelial cells (HUVECs) on Matrigel and inducing tube formation with vascular endothelial growth factor (VEGF). HUVECs in the basal state where no VEGF is added exhibit limited angiogenesis, forming short connections and minimal junctions (**Figure 1**). The addition of VEGF (15 ng/mL) dramatically increases the sprouting length and number of junctions creating an extensive capillary-like network after 6 hours at 37°C. Treatment with 1 nM of BIIG2 or MINT1526A was unable to disrupt the VEGF-stimulated tube formation as the segment lengths and the number of tube junctions are minimally decreased (**Figure 1**). BIIG2 at 10 nM and 100 nM reduced both the total segment lengths and the number of junctions whereas MINT1526A only reduced segment lengths at 100 nM. MINT1526A showed a trend towards reducing the number of tube junctions but did not reach statistical significance (P < 0.05). Of the concentrations tested, BIIG2 reduces both branching and extension capabilities of HUVECs in a dose-dependent manner. In contrast, MINT1526A only disrupts extension and branching is maintained. Based on these results, we sought to further investigate the inhibitory effects of angiogenesis in three dimensions using HUVEC spheroids (**Figure S2**). HUVECs were embedded into a collagen gel and sprouting was assessed 24 hours after being treated with 25 ng/mL VEGF. Sunitinib, a tyrosine kinase inhibitor known to inhibit angiogenesis ^26^, was used as a positive control and reduced VEGF-induced HUVEC sprouting in a dose-dependent manner. Unlike the two-dimensional tube formation assay, BIIG2 and MINT1526A did not noticeably reduce spheroid sprouting.

**Figure 1.**
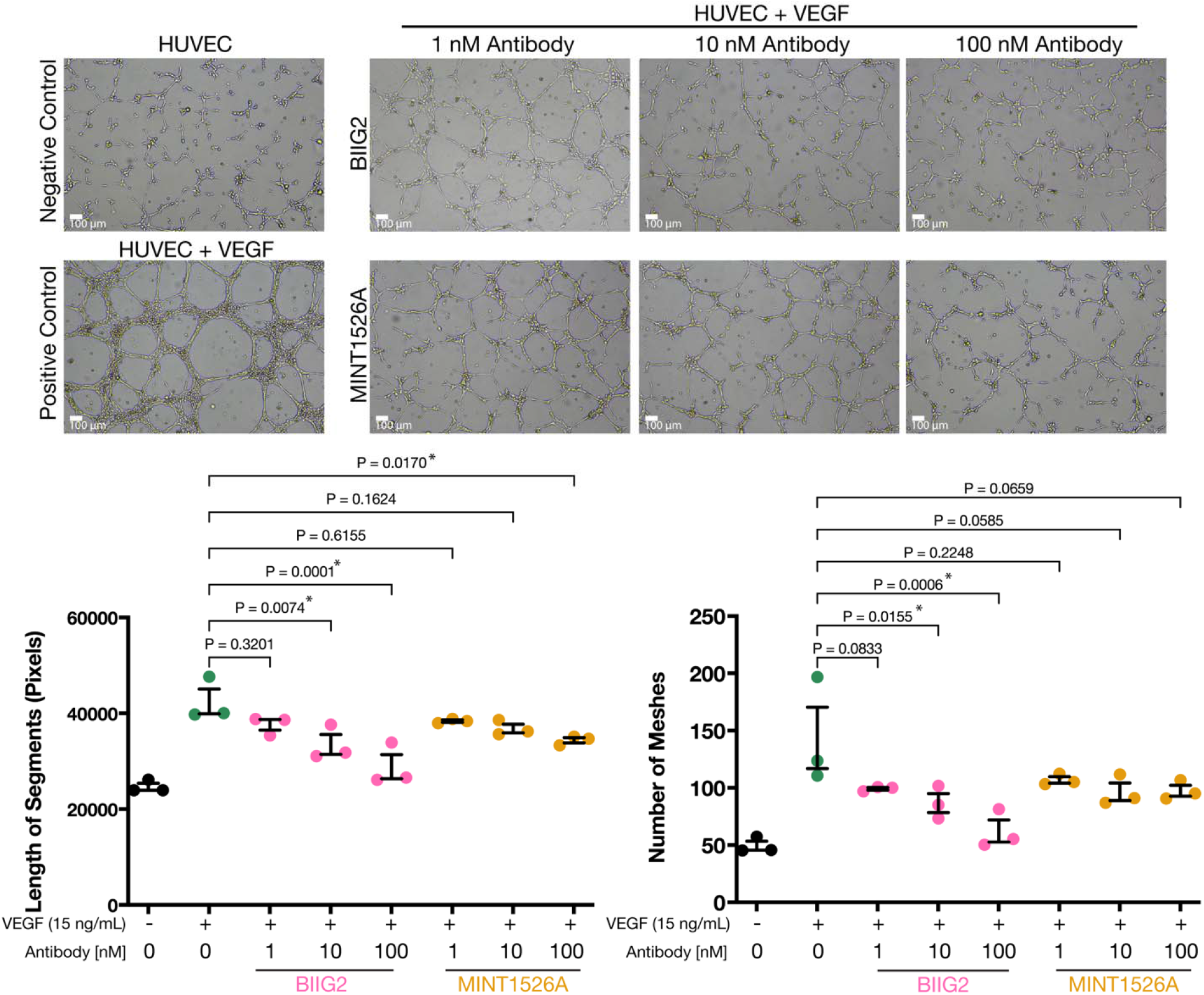
Effects of BIIG2 and MINT1526A α5 integrin antibodies on endothelial tube formation *in vitro*. Representative images of HUVEC tube formation by VEGF on Matrigel after 24 hours of incubation in the presence or absence of antibodies. Scale bar is 100 μm. HUVEC cultured in medium alone served as the basal control, whereas HUVEC treated with VEGF (15 ng/mL) were used as the positive control. The quantification of tube formation involved measuring the total length of master segments (the cumulative length of tubes formed, measured in pixels) and the number of meshes using angiogenesis analyzer for ImageJ ^81^. Master segments and mesh counts were averaged from three fields of view per replicate, n=3 (9 total images, 3 images per replicate). Data are shown as the mean ± SEM; n=3. One-way ANOVA with Bonferroni correction; adjusted P values are shown.

### BIIG2 localizes to connective tissue-resident fibroblasts and melanoma cells

Integrin α5β1 is expressed in various tissues at low levels ^19^. We used immunohistochemistry to determine the cellular localization and tissue specificity of BIIG2 and compared the polyclonal α5 integrin antibody PA5-82027. As expected, PA5-82027 stains a broad spectrum of human tissues, including placental syncytiotrophoblasts, striated ducts in salivary glands, and alveolar ducts (**Figure S3A and Table S1**). In contrast, BIIG2 exhibits a restricted staining profile, predominantly marking isolated connective tissue-resident fibroblasts across various tissues. Remarkably, BIIG2 also demonstrated intense staining of activated and transformed melanocytes in the epidermis and upper dermis derived from patients with atypical nevi and invasive cutaneous melanoma (**Figure S3B**). Tissue specificity in therapeutics is advantageous since it has the potential to reduce off-target effects and BIIG2’s specificity supports the targeted treatment of various pathologies, including fibrosis, sclerosis, and melanoma.

### BIIG2 reduces tumor volume in a patient-derived xenograft model

Based on the strong BIIG2 immunohistochemical staining localized to human melanoma cells (**Figure S3B**), and considering the known tumor-inhibitory effects of other anti-α5 antibodies in rhabdomyosarcoma and glioblastoma mouse models ^27,28^, we established two melanoma xenograft cultures to evaluate the *in vivo* inhibitory effects of BIIG2 on melanoma growth. Using brain metastases from patients who had different responses to treatment, we derived two cultures, M15 and M12 (**Figure S4A-B**). M15 cells contained the *NRAS^Q^*^61^ mutation and exhibited metastatic characteristics and resistance to radiation and systemic therapy. M12 cells contain the *BRAF^V^*^600^ mutant and responded well to systemic therapy with the donor patient achieving complete remission ^29^. Quantitative PCR of M15 and M12 shows differing levels of melanoma markers compared to normal human melanocytes (NHM) such as melan-A (*MLANA)* and microphthalmia-associated transcription factor (*MITF)*, *CDKN2A* tumor suppressor as previously described ^30–32^. M15 cells have a higher mRNA expression of extracellular matrix proteins such as: fibronectin *(FN1)* and osteopontin (*SPP1)*, but lower for tenascin C (*TNC)* compared to M12 (**Figure S4C**). Furthermore, cell surface integrin flow cytometry showed that compared to NHM, αv and α5 are upregulated in M15 and M12 cells, with the highest α5 expression in M15 cells agreeing with previous reports (**Figure S4D**) ^17,33^.

M15 and M12 cells were embedded in Matrigel to grow and subsequently injected into the flanks of athymic mice to determine the effect on tumor volumes *in vivo* (**Figure 2A**). When tumors became palpable, antibodies were administered intraperitoneally at a dosage of 10 mg/kg twice weekly. Treatments were stopped when the tumors injected with control IgG surpassed an average volume of 1000 mm^3^. In the less aggressive M12 xenograft, BIIG2 demonstrates a measurable anti-tumor effect (**Figure 2B**). However, it is not more effective than the P1D6 antibody or antibodies targeting integrin αv (**Figure S5A**). Expression analysis revealed that in BIIG2-treated mice, the xenografted M12 tumors exhibited an increase in RNA levels of the melanocyte differentiation marker *MLANA* and *p53* by 50% and 32% respectively, in comparison to those treated with control IgG (n = 9 mice per group; T-test p < 0.01; **Figure S5B**). In the more aggressive M15 tumor challenge model, no antibodies tested demonstrated a statistically significant impact on tumor growth. In some replicates, BIIG2 demonstrated clear inhibitory effect, however the overall mean did not reach statistical significance. Pentamidine isethionate (Pentam) has been shown to restore free p53 levels in melanoma cells, inhibiting tumor growth and combination targeted therapies have shown improvement and promise for melanoma treatment ^34,35,36^. We found this was the case for M15 treatment where neither Pentam nor BIIG2 alone reduced tumor volume, however, when combined, a significant reduction of tumor volume was achieved (**Figure 2C**). In M12, Pentam and BIIG2 were both effective in reducing tumor volumes as monotherapies and their combination demonstrated efficacy, but was not more effective. The clinically tested MINT1526A could not be used for comparison due to a shortage of purified antibody, which was kindly provided by Genentech.

**Figure 2.**
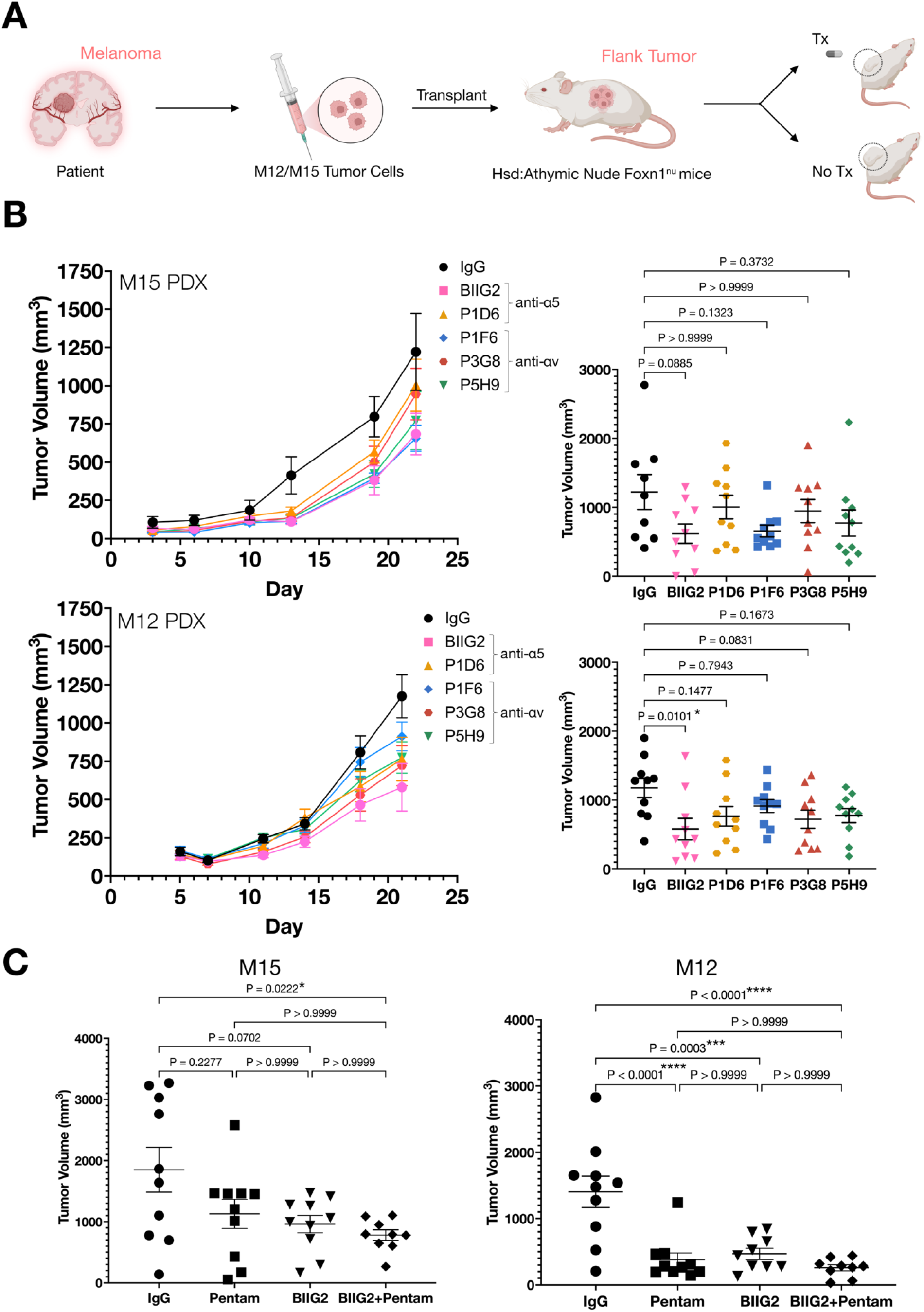
*In vivo* anti-tumor activity of BIIG2. **(A)** Athymic nude mice were xenografted with patient-derived melanoma cells (labeled M15 and M12) from brain metastasis. Upon observation of palpable tumors, we proceeded with the treatment of the mice. Illustration created with BioRender.com. **(B)** Mice were injected intraperitoneally with BIIG2 or other αv and α5 integrin antibodies. Treatment was administered twice weekly at a dose of 10 mg/kg. Data plotted are shown as means ± SEM; n=10 per day of measurement. Tumor volumes on the final day of observation were plotted as mean ± SEM; n=10, except for M15 where IgG control n=9. A one-way ANOVA with Bonferroni correction was performed for statistical significance of the complete antibody panel against the control IgG, adjusted P values are shown. Comparisons between BIIG2 and the panel of antibodies were analyzed in the same method in Figure 5A. **(C)** Pentamidine isethionate, an aromatic diamidine with anti-cancer activity, was administered intraperitoneally at a dose of 10 mg/kg once daily. Data shown are mean ± SEM; n=10 mice per treatment group; one-way ANOVA with Bonferroni correction; adjusted P values are shown. The clinically tested MINT1526A could not be used for comparison due to a shortage of purified antibody, which was kindly provided by Genentech. See also Figures S4 and S5.

### BIIG2 and MINT1526A bind α5β1 with sub-nanomolar affinity independent of cation condition

Having established the potential of BIIG2 in angiogenesis and melanoma tumor growth inhibition, we characterized the affinities of BIIG2 and MINT1526A to α5β1, a critical consideration for effective therapeutics ^37^. Integrin function, signaling, and its ability to bind ligands are linked to its conformation and can be influenced *in vitro* using divalent cations ^1,38^. In β1 integrins, calcium (Ca^2+^) is known to promote a bent conformation which is linked to inactive integrins, while manganese (Mn^2+^) favors an open conformation, which is linked to integrin activity ^39–41^. Using recombinant human α5β1 ectodomain (**Figure S1A**) in the presence of 5 mM Ca^2+^ or 1 mM Mn^2+^, we employed biolayer interferometry (BLI) to determine if the conformation of α5β1 impacts the ability of BIIG2 or MINT1526A to bind. We find both antibodies bind α5β1 in the low nanomolar range irrespective of cations: BIIG2 binds to α5β1 with an affinity of 0.57 nM in Ca^2+^ and 0.64 nM in Mn^2+^; MINT1526A binds α5β1 with an affinity of 1.3 nM in Ca^2+^ and 0.23 nM in Mn^2+^, thus demonstrating that regardless of cation condition, both antibodies bind (**Figure 3A**). Despite these similarities, the antibodies have distinct profiles: in both conditions, BIIG2 associates (*K_on_*) and dissociates (*K_off_*) slightly more slowly than MINT1526A, the slower *K_off_* suggesting better stability. Although for BIIG2, *K_D_* is similar in both cation conditions, for MINT1526A, there is a marked increase in affinity in the presence of Mn^2+^ vs. Ca^2+^, suggesting Mn^2+^ stabilizes binding (**Figure 3A**).

**Figure 3.**
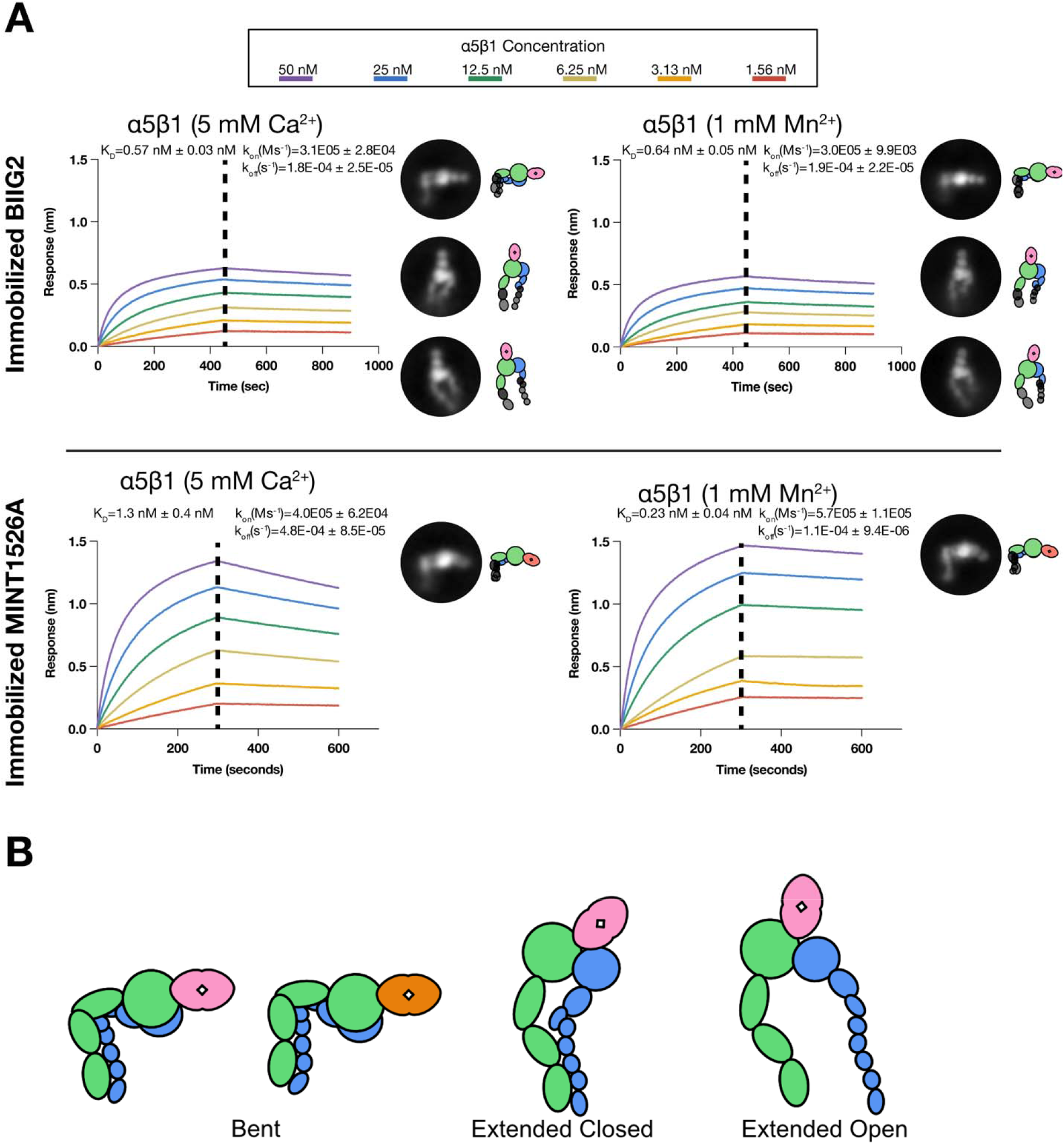
Biolayer interferometry binding kinetic analysis and negative stain electron microscopy of integrin-Fab complexes. **(A)** Representative BLI sensorgrams displaying the association and dissociation step of the binding assay between immobilized BIIG2 and MINT1526A and α5β1 integrin ectodomain in the presence of Ca^2+^ (nonactivating) and Mn^2+^ (activating). BIIG2 binds α5β1 ectodomain with a *K*_D_ of 0.57 nM and 0.64 nM in nonactivating conditions and activating conditions respectively. MINT1526A binds α5β1 ectodomain with a *K*_D_ of 1.3 nM and 0.23 nM in nonactivating conditions and activating conditions, respectively. Each experimental condition was repeated three times with the K_D_, *k_on_* (1/Ms) and *k_off_* (1/s) reported as the mean ± SEM. Representative negative stain EM 2D class averages of the α5β1 integrin in complex with the antibody Fab fragment in their respective buffer conditions. The complete set of 2D class averages are in **Figure S7A**. **(B)** A cartoon schematic representing the conformations of integrin observed under negative stain EM with either BIIG2 or MINT1526A Fab bound. Pink, BIIG2; Orange, MINT1526A. Binding of BIIG2 Fab does not influence the conformational landscape of α5β1. MINT1526A-bound α5β1 is only observed in the half-bent conformation. See also Figure S6.

We hypothesized that the antibody-mediated reduction in angiogenesis is due to steric inhibition of α5β1 binding to fibronectin. To test this hypothesis, we conducted a series of competition experiments using BLI to determine if BIIG2 and MINT1526A prevented α5β1 from binding to fibronectin and if fibronectin could prevent BIIG2 or MINT1526A binding (**Figure S6**). We used a recombinant truncated fibronectin fragment (FN7-10) that contains the necessary domains for α5β1 binding ^12,15,42^ and consistent with prior studies, we show that α5β1 binds FN7-10 with a high affinity in the presence of Mn^2+^, and Ca^2+^ significantly reduces the affinity and binding signal ^12,39^. In the presence of Mn^2+^, we find that if α5β1 is preincubated with either BIIG2 or MINT1526A, there is a marked decrease in binding signal to immobilized FN7-10. Likewise, if α5β1 is preincubated FN7-10, both antibodies do not show binding, suggesting that FN7-10 is unable to displace bound antibody and antibody is unable to displace bound FN7-10. When alone, FN7-10 binds α5β1 with an affinity of 2.2 nM, but if α5β1 has been preincubated with BIIG2 FN7-10 binds with 9.2 nM affinity and if preincubated with MINT1526A the affinity of FN7-10 is even lower at 13.8 nM. Likewise, after preincubating integrin with FN7-10, subsequent antibody binding was drastically reduced: immobilized BIIG2 bound α5β1 with 0.689 nM affinity, but if α5β1 was preincubated with FN7-10, this dropped to 14.6 nM. The effect on MINT1526A was most drastic, dropping from 0.18 nM to 25.1 nM. This suggests that the antibodies, although capable of binding α5β1 with extremely high affinities, are unable to displace FN7-10 once it is bound. Taken together, this suggests that the fibronectin, BIIG2 and MINT1526A share a similar or overlapping binding interface. As anticipated, α5β1 does not bind FN7-10 with high affinity in high Ca^2+^ condition and thus we show that in high Ca^2+^, BIIG2 and MINT1526A bind integrin with only slightly lower affinity than if FN7-10 is not present.

### MINT1526A preferentially binds compact α5β1; BIIG2 binds multiple conformations

To visualize epitopes and the impact of antibody binding on α5β1’s conformation, we used single-particle negative stain electron microscopy (nsEM) to visualize the protein complexes of purified α5β1 ectodomains bound to BIIG2 or MINT1526A Fabs. We generated 2D class averages that summed thousands of individual particle images (**Figure S7A**). As previously hypothesized, our nsEM analysis confirmed that both BIIG2 and MINT1526A bind the α5 subunit and we further localized the epitope to the α5 β-propeller domain (**Figure S1A**) ^22,23^. Fibronectin binds the same domain, which further supports the hypothesis that both antibodies inhibit α5β1 function by blocking fibronectin binding through steric hindrance.

In nonactivating conditions (5 mM Ca^2+^), most (98%) α5β1 adopts two conformations: ‘bent’ and ‘extended closed’ (**Figure S7A**), consistent with previous work ^39^. In activating conditions (1 mM Mn^2+^), we observed an increase in the proportion of α5β1 particles adopting the’extended open’ conformation (24% vs 2%). When α5β1 is bound to the BIIG2 Fab, the ratios of integrins in the three different conformations are similar in both cation conditions (**Figures 3A and S7A**). In contrast, when MINT1526A binds, we only observe the bent conformation in both activating and nonactivating conditions (**Figure S7A**). We considered two possibilities: either MINT1526A binds any integrin conformation and then “locks” it into a bent conformation or it binds only the bent conformation (which is present in both cation conditions due to the dynamic equilibrium) and then locks it. To address this point, we incubated α5β1 with the antibody 9EG7 to induce an extended conformation and then added MINT1526A ^43^. The resulting nsEM did not yield any 2D class averages with both 9EG7 and MINT1526A bound, suggesting MINT1526A has a strong preference for the bent conformation. In contrast, when we conducted a similar experiment using BIIG2 we found approximately 33% of particles had both 9EG7 and BIIG2 bound (**Figure S7B**).

In summary, although both antibodies inhibit α5β1 function by blocking fibronectin binding through steric hindrance, MINT1526A has an additional conformational effect, restricting the observed conformational equilibrium and preventing integrin extension and headpiece opening.

### High-resolution structural analysis of the BIIG2 Fab reveals occlusion of the integrin binding cleft

We determined the structure of the BIIG2 Fab to 1.4 Å resolution using X-ray crystallography (PDB ID: 8R38; **Figures 4F and S8K-M**). The structure was refined to *R*/*R*_free_ values of 19.9/22.4% with good geometry (**Table S2**), and is characterized by well-defined electron density from residues 1-210 and 1-214 (Kabat numbering) of the heavy and light chains, respectively. In particular, the antigen-binding site is well-defined. The binding site of the BIIG2 light chain is dominated by positive charges from three lysine residues (K27, K31, and K92) and R28, whereas the heavy chain CDRs are charge-balanced. The BIIG2 CDR H3 loop is exposed, notably longer than the other CDRs and partially involved in crystal contacts through residues T97, M98, G99, and CDR L1 loop R27 (**Figure 4F**). The obtained high resolution allowed for accurate modeling of several modified residues. These include a pyroglutamate on the N-terminal position of the heavy chain, a methyllysine on the light chain (L199), and a 5-hydroxyproline modification in the CDR H3 loop (Pro 100C). In addition, we observed well-defined electron density for the only visible glycosylation on BIIG2, located on glycan N60 of the Fv heavy chain.

**Figure 4.**
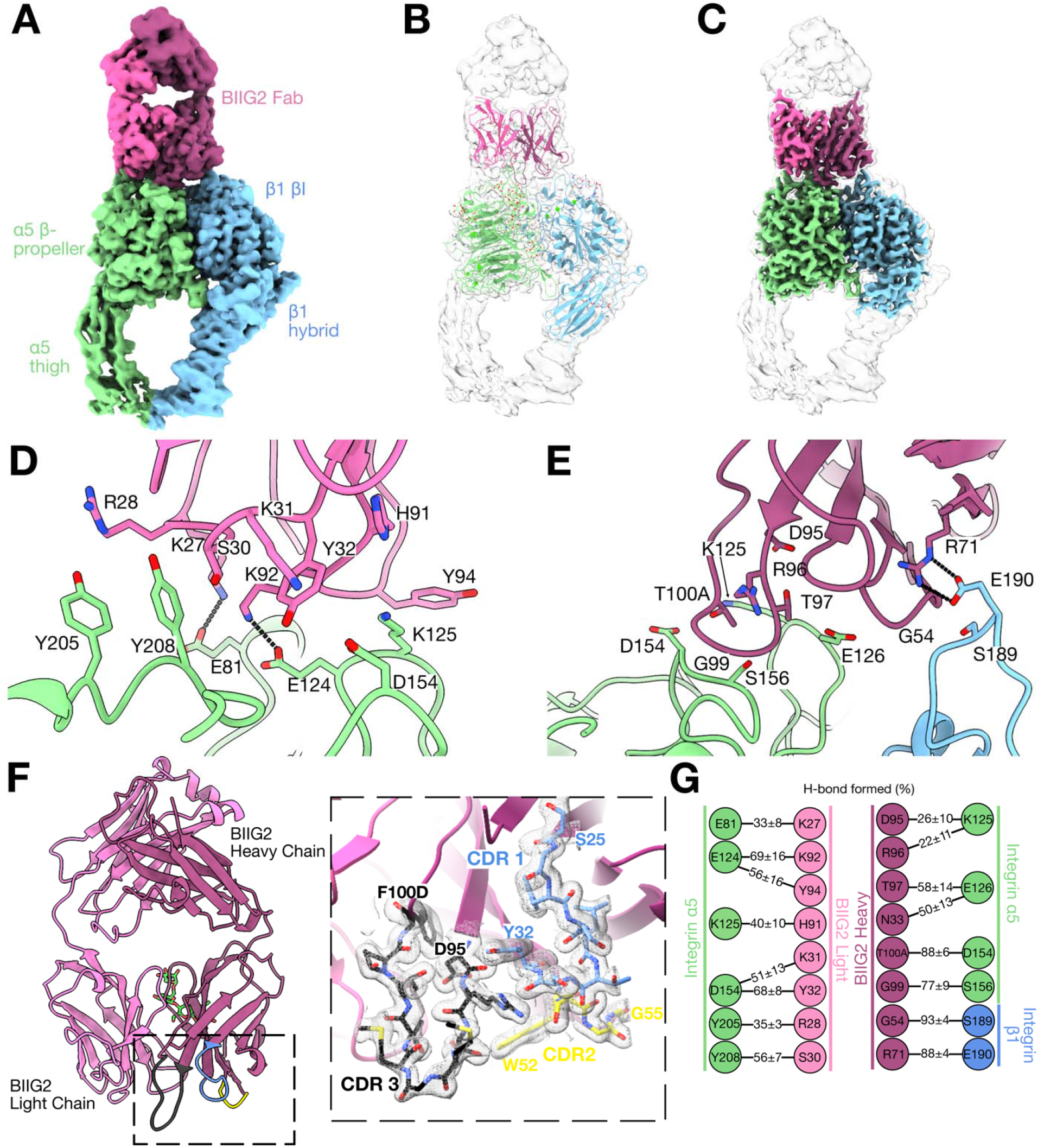
CryoEM analysis of integrin α5β1:BIIG2 Fab complex. **(A-C)** CryoEM maps and model of the α5β1 ectodomain with BIIG2 Fab bound. The color code is as follows: integrin α5 subunit is green, integrin β1 subunit is blue, BIIG2 Fab light chain is light pink and heavy chain is dark pink. **(A)** The unsharpened map at a low threshold. **(B)** A ribbon diagram (colors) in the cryoEM map (semi-transparent white). **(C)** A higher threshold sharpened cryoEM map (colors), superimposed with the unsharpened map (semi-transparent white). **(D and E)** A close-up view of the interface between α5β1 and the light chain **(D)** and heavy chain **(E)** of the BIIG2. Salt bridges are drawn with dashed lines. **(F)** The crystal structure of BIIG2 Fab (light chain in light pink, heavy chain in dark pink). The single N-linked glycan N60 is shown as green stick representation. A close-up view of the Fab crystal structure shown in cartoon representation (dashed box), with σ_A_-weighted 2*F*_o_-*F*_c_ electron density map (grey mesh, contoured at 1.0 σ) for selected residues of the CDR loops (CDR1-yellow, CDR2-blue, and CDR3-gray), which are depicted in stick representation; amino acid residues are labeled. **(G)** Percentage of hydrogen bonds and salt bridges formed (mean ± SEM) between integrin α5β1 and BIIG2 Fab molecular dynamics simulations using a 3.5 Å cutoff between acceptor and donor-H atoms. Values were averaged across seven simulation repeats, each 2.0 µs long. Stable interactions are summarized in **Table S4.** See also Figures S8 and S10.

To map how BIIG2 binds to and inhibits α5β1 function, we used single-particle cryogenic electron microscopy (cryoEM) to determine the structure of the α5β1:BIIG2 Fab complex to a global resolution of 2.7 Å (**Figure 4A and Table S3**). After focused refinement, the α5β1 headpiece/BIIG2 Fab variable-region interface was estimated at 2.5 Å resolution, which allows for modeling (**Figures 4B-C, S8 and S10**). The crystal structure of the BIIG2 Fab (PDB ID: 8R38; this work) and the close α5β1 closed headpiece structure (PDB ID: 3VI3; ^44^) were docked (rigid body fitting) into our cryoEM density as starting coordinates for model building. From here, we built the α5 β-propeller, the β1 β-I domain and a portion of the hybrid domain, and the variable region of the BIIG2 Fab. As with many cryoEM maps of integrins ^12,45–50^, our map is anisotropic, with the resolution tapering off away from the well-resolved headpiece. Secondary structural elements for the α5 thigh and β1 hybrid domains are visible, but the resolution is insufficient for residue-level modeling. Additional 3D classification reveals populations of BIIG2-bound integrin in the bent (25%) and extended closed conformations (18%). We note that a significant number of particles (57%) contribute to classes that do not have a strong density for the lower leg region (**Figure S11B**). We hypothesize that the lower legs could be averaged out due to significant conformational variability or interactions with the air-water interface that compromise the particle integrity and note that all domains are visible in nsEM averages and the molecular weight estimate from size exclusion chromatography (SEC) is consistent with a full α5β1 ectodomain (**Figures S7 and S8**).

The cryoEM map reveals that BIIG2 engages both the α5 and β1 integrin subunits through electrostatic interactions, with positively charged residues on BIIG2 and key negatively charged residues on the integrin. Unlike fibronectin, BIIG2 does not contain an RGD motif nor does it bind to any RGD-interacting residues on α5β1. Two lysine residues on the BIIG2 light chain (K27 and K92) form salt bridges with the known negatively charged α5 synergy site residues E81 and E124 (**Figure 4D**). This portion of the epitope on the α5 β-propeller is further stabilized by surrounding hydrogen bonds. BIIG2 engages several residues that are necessary for efficient FN binding: D154, a key residue stabilizing the interaction between α5 and the FN9 synergy site, interacts with T100A of the BIIG2 heavy chain, thus occluding FN9 access to this residue (**Figure 4E**) ^12,44,51^. Additionally, R71 of the BIIG2 heavy chain forms a salt bridge with E190 of the Specificity-Determining Loop 2 (SDL2) of the β1 subunit, providing additional stabilizing interactions in the α5β1:BIIG2 complex (**Figure 4E**). BIIG2’s CDR H3 methionine residues M98 and M100 exhibit long-range sulfur-aromatic interactions with α5 W157 and F155 respectively, further stabilizing the binding interface (**Figure S12D**) ^52^. To further support these observations, we employed all-atom molecular dynamics (MD) simulations of the α5β1 headpiece complex with the BIIG2 Fab (**Figure S12A-C**). Two microsecond-long trajectories show that the network of hydrogen bonding and electrostatic interactions observed in the cryoEM structure are dynamically involved in maintaining the α5β1:BIIG2 interface (**Figure 4G**). These interactions, summarized in **Table S4**, position BIIG2 to sterically occlude fibronectin from accessing the primary RGD binding site and α5’s synergy binding site for FN9 (**Figure S13A**).

### MINT1526A exclusively binds the α5 subunit and prevents fibronectin binding by blocking the synergy binding site

To directly compare the binding interfaces with the BIIG2 complex, we next determined the cryoEM structure of the MINT1526A Fab bound to the α5β1 ectodomain to a global resolution of 3.1 Å (**Figure 5A and Table S3**). Similar to the α5β1:BIIG2 Fab cryoEM map, the resolution of α5β1:MINT1526A Fab was anisotropic, with the highest-resolution regions (2.8 Å) in the headpiece and at the epitope/paratope interface (**Figures 5C, S9 and S10**). 3D classification reveals that the integrin angle of rotation about the knee region ranges from 90 to 73 degrees, in agreement with previous data (**Figure S11D**) ^12^. MINT1526A exclusively binds the α5 subunit and it does not block the canonical RGD binding site cleft. Although MINT1526A also does not directly interact with any residues on α5β1 required to mediate FN binding, its epitope overlaps with one of the known FN binding sites, preventing interactions between α5 and the FN9 synergy site (**Figure S13B;** ^12,51^). α5’s would-be D154 electrostatic and Y208 proline-aromatic interaction with FN are sterically blocked by the CDR H3 of MINT1526A (**Figure 5E**). The high affinity of α5β1:MINT1526A is achieved primarily by hydrogen bonds, involving residues throughout the epitope (**Figure 5E-F**). MD simulations of α5β1:MINT1526A Fab indicate that these hydrogen bonding interactions between MINT1526A and α5β1 are stable (**Figure 5D**). A summary of the interactions can be found in **Table S4.**

**Figure 5.**
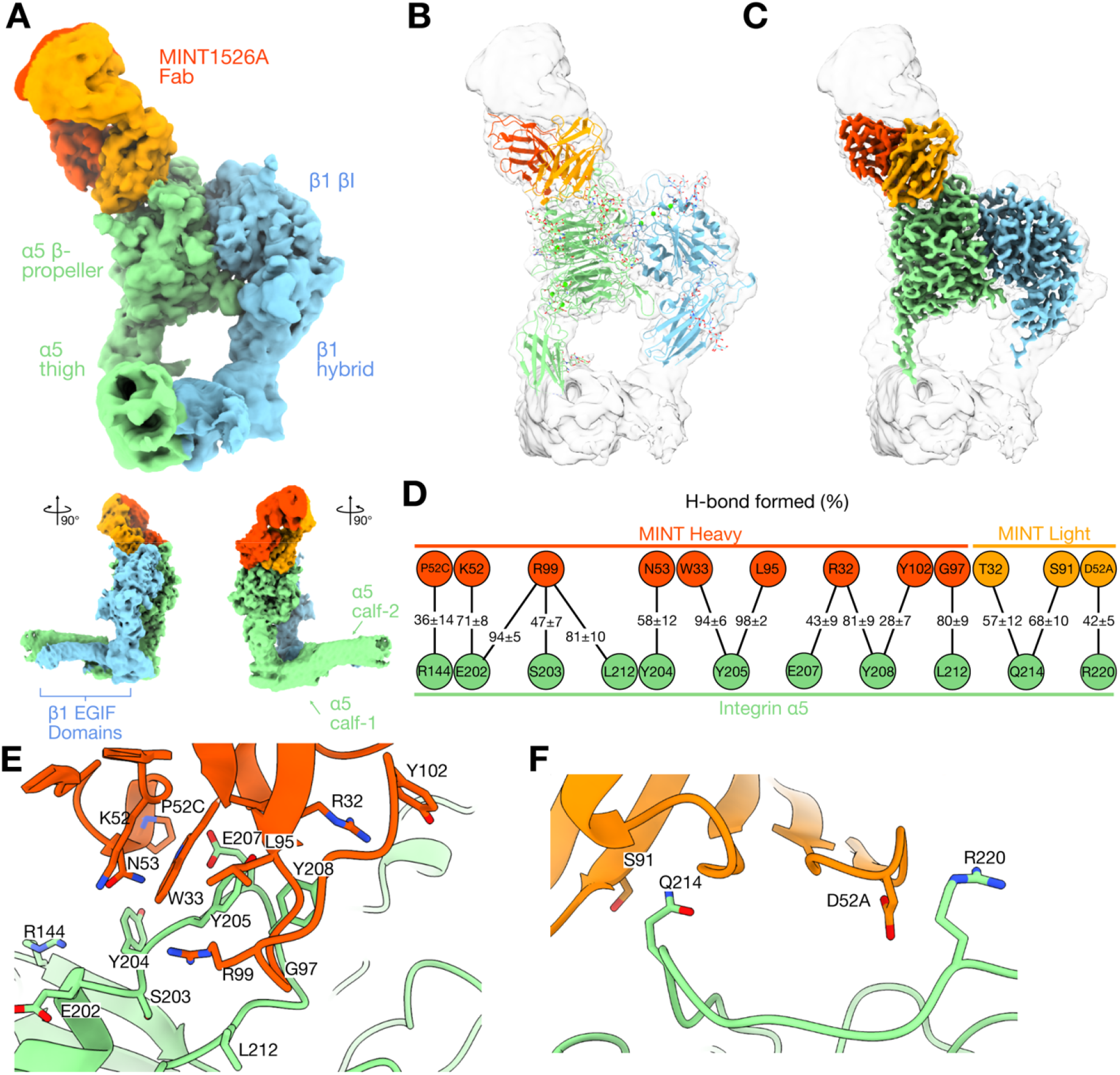
CryoEM analysis of α5β1:MINT1526A Fab complex. **(A-C)** CryoEM maps and model of the α5β1 ectodomain with MINT1526A Fab bound. The color code is as follows: integrin α5 subunit is green, integrin β1 subunit is blue, MINT1526A Fab light chain is light orange and heavy chain is dark orange. **(A)** The unsharpened map at a low threshold. **(B)** A ribbon diagram (colors) in the cryoEM map (semi-transparent white). **(C)** A higher threshold sharpened cryoEM map (colors), superimposed with the unsharpened map (semi-transparent white). **(D)** Hydrogen bonds identified between integrin α5β1 and MINT1526A Fab are supported by molecular dynamics simulations. Percentage of hydrogen bonds formed (mean ± SEM) between integrin α5β1 and MINT1526A Fab using a 3.5 Å cutoff between acceptor and donor-H atoms. Values were averaged across seven simulation repeats, each 2.0 µs long. A ribbon diagram shows a close-up view of the interface between α5β1 and the MINT1526A Fab heavy chain **(E)** and light chain **(F)**. Stable interactions are summarized in **Table S4.** See also Figures S9 and S10.

## Discussion

Integrins play an essential role in cell proliferation, adhesion, migration, and survival, and their dysregulation is attributed to the progression of numerous human diseases including cancer ^4^. Due to its important role in tumor angiogenesis, α5β1 has long been identified as a target for cancer therapies ^4,19,53^. More recently, antibodies targeting α5 integrin have been demonstrated to inhibit the progression of the neurodegenerative disorder amyotrophic lateral sclerosis (ALS) and idiopathic pulmonary fibrosis (IPF), underlining and expanding α5β1’s potential as a therapeutic target ^54,55^. Understanding the precise molecular interactions mediating integrin-therapeutic interactions will be key for the development of tailored, specific, and effective therapeutics, however, the molecular details of α5β1 with any function-modulating antibody have so far been lacking. We present a comprehensive functional, structural, and mechanistic characterization of BIIG2, a rat IgG antibody targeting human α5 integrin, demonstrating properties that support further therapeutic evaluation. BIIG2 exhibits high affinity for α5β1 and shows intense, tissue-specific binding to connective fibroblasts and melanocytes in melanoma patient tissues, which are known to express elevated levels of active α5β1 integrin. This staining pattern may result from BIIG2’s ability to recognize the full spectrum of α5β1 headpiece conformations. BIIG2 more effectively reduces VEGF-induced endothelial tube formation compared to MINT1526A (RG-7594), an anti-α5β1 antibody that has already been tested in clinical trials. In the M12 patient-derived xenograft mouse model, BIIG2 demonstrates reduced tumor growth relative to other anti-integrin antibodies, with efficacy comparable to p53 restoration via pentamidine isethionate.

Using single-particle cryoEM and BLI, we show that both BIIG2 and MINT1526A have similar affinities but different kinetics for α5β1, and impede activation by sterically blocking fibronectin binding, albeit through different epitopes. These kinetic differences suggest that the two antibodies engage α5β1 through distinct binding pathways. MINT1526A associates more rapidly under both cation conditions, whereas dissociation depends on cation condition. BIIG2 engages both the α5 β-propeller and SDL2, while MINT1526A contacts only the α5 β-propeller, which may contribute to their different kinetic and conformational profiles. Alternatively this could be because whereas BIIG2 binds most integrin conformations, MINT1526A is more selective. This is consistent with our negative stain analysis, suggesting that MINT1526A has a significant preference for binding bent integrin (**Figure S7**). It is intriguing to note, however, that despite preferentially binding the bent conformation, MINT1526A is stabilized in the presence of Mn^2+^, the cation known to stabilize the active, extended conformation.

The dramatic difference in the available conformational states of BIIG2-bound α5β1 vs MINT1526A-bound α5β1 was unexpected, since neither Fab interacts with residues that are known to participate in integrin α5β1 extension or headpiece opening ^11,12,43,44,56^. The root-mean-square difference (RMSD) of the α5β1 headpiece region in these two models is 0.7 Å, indicating that there are no significant global structural differences when BIIG2 or MINT1526A is bound. The most notable difference in our two integrin models is a loop (residues 27-33) between two β-sheets in the first blade of the α subunit’s β-propeller. It is tempting to speculate that like other flexible loops in the α subunit ^57^, this positioning could be related to the overall conformation of α5β1. In the MINT1526A-bound bent α5β1 structure, the loop points downward, similar to the recent cryoEM structure of α5β1 in a bent conformation (PDB ID: 7NXD), whereas the BIIG2-bound α5β1 has loop points upward, similar to an extended structure from the same study bound to fibronectin (PDB ID: 7NWL; ^12^). Most published structures of this loop are more similar to the ‘up’ position of the loop, however, since many of these structures are truncated integrin fragments that do not include the lower leg domains, we cannot be certain if this loop is related to overall conformation ^44,56,58^. Therefore, the underlying cause of the observed differences in α5β1 conformations is not entirely clear although allosteric changes must either be transmitted or recognized by MINT1526A.

Although integrins have long been identified as promising therapeutic targets, with over 130 clinical trials investigating integrin-targeting therapies, fewer than a dozen therapeutics– either antibodies or small molecules– have reached the market ^59^. Several cancer-related integrin therapeutics have progressed to late-stage clinical trials although none have been approved to date ^60^. Volociximab (M200) was well tolerated in phase 1 trials ^61^, but demonstrated limited efficacy in phase 2 across multiple cancer types leading to discontinuation of further development ^20,62^. Many other investigational integrin-targeting therapies in oncology have also been halted due to limited efficacy or safety concerns. Nevertheless, integrins remain validated therapeutic targets, with several approved agents demonstrating significant success in non-oncologic indications ^7^. Interest in the field remains strong, with numerous ongoing trials showing promising early results.

One hypothesis posits that the success or failure of some integrin therapeutics depends on selectively targeting specific integrin conformations. This concept is exemplified by Nimotuzumab, an anti-tumor antibody targeting the epidermal growth factor receptor (EGFR). Nimotuzumab permits EGFR to adopt its active conformation and exhibits fewer side effects than other anti-EGFR antibodies, such as Cetuximab, which bind similar epitopes but do not allow the active receptor conformation ^63^. For integrins, the selectivity of therapeutics for specific conformations or their influence on integrin conformation, and how those differences translate into tissue specificity, off-target effects, or toxicity remains incompletely characterized, although recent work has begun to shed light on these mechanisms ^50,58,64,65^. Structural and biochemical data are available for several late-stage and marketed therapeutics, including high-resolution structures of the epitope or binding site for five of the seven approved integrin therapeutics ^48,64,66–69^. These therapeutics also target the headpiece region near or in the ligand-binding cleft with steric clash with the cognate integrin ligand as the proposed mechanism, similar to what our structures of BIIG2 and MINT1526A show. Of these complexes, the four solved by crystallography show a single conformation, and of the three determined via cryoEM, abciximab:αIIbβ3 shows a closed headpiece, eptifibatide:αIIbβ3 shows an open headpiece, M-tirofiban:αIIbβ3 is in an acutely bent conformation. The only cryoEM structure showing heterogeneity is tirofiban:αIIbβ3 showing an open headpiece with heterogeneity in the α leg ^48,66,69^. Thus, the conformational landscape available to the therapeutic-bound integrins remains incomplete ^7^.

Recently, it has been proposed that partial agonism – linked to stabilizing the open conformation – could be responsible for chronic indications of certain failed integrin therapeutics. A comprehensive study of αIIbβ3 small molecule inhibitors showed that of the inhibitors tested, stabilizing the bent conformation has led to better therapeutic outcomes than those that stabilize an open conformation. That study also identified a general chemical principle for the design of bent-stabilizing inhibitors and, excitingly, extended the work to show the principle held true for the distantly related integrin α4β1 ^64^.

On the other hand, studies show that small molecule and antibody inhibitors that stabilize the bent-closed conformation of αvβ6 are not able to lower expression of fibrotic-associated genes in lung epithelial cells as effectively as inhibitors that stabilize the extended-open conformation ^70^. Clinical advancement of two potential therapeutics that stabilized the closed αvβ6 conformation, antibody BG00011 (the humanized version of 3G9) and the small molecule MORF-720 were halted due to suspected on-target safety signals ^55,71,72^. In contrast, PLN-74809, which binds all conformations of αvβ6 and αvβ1 and GSK3008348, which causes rapid internalization of αvβ6 which is associated with an active conformation, have led to more promising initial results with no serious adverse effects related to the compounds in clinical trials ^73–76^. Similarly, Etaracizumab, an enhanced form of Vitaxin, is well-tolerated in patients and currently under investigation for cancers ^77,78^. Its precursor, LM609 has been shown to bind all conformations of integrin αvβ3 ^79^.

It has become clear that the challenges associated with integrin-therapeutic development remain numerous, however, exciting recent results linking functional and known clinical outcomes to structural interactions have enabled recent leaps in the establishing of principles that govern effective therapeutics^64^. Significant efforts have advanced the discovery of novel integrin therapeutics including a new platform for antibody discovery ^80^ and the development of *de novo* structure-based rationally designed integrin miniproteins ^50,58^. In addition to the specific structural and functional details of α5β1-antibody interactions, the results presented here contribute to the greater understanding of the complex interplay between potential integrin therapeutics, conformation, and downstream effects.

### Limitations

While this study advances our understanding of the dynamics of therapeutic antibodies and their impact on integrin conformation equilibrium, it has limitations. For a more complete comparison of efficacy to BIIG2, the xenograft tumor model data using MINT1526A would be needed. While we are able to make a direct comparison within our i*n vitro* experiments, we are not able to repeat this for our xenograft experiments due to insufficient quantities of antibody available. Additionally, all structural experiments were conducted using integrin ectodomain constructs and it is possible that full-length integrins (containing the transmembrane and cytoplasmic domains) could show different conformational dynamics. Finally, although we performed a biophysical analysis, an in-depth nsEM study using the native ligand and a conformational antibody, and resolved a high-resolution cryoEM map of the α5β1-MINT1526A complex, we were unable to identify the specific allosteric molecular details that lead to the bent-conformation binding preference or the headpiece “locking” we observe.

## Resource Availability

### Lead contact

Further information and request for resources and reagents should be directed to and will be fulfilled by the lead contact, Melody G. Campbell.

### Materials availability

All plasmids generated in this study are available from the lead contact.

### Data and code availability

All structures and models referenced have been deposited in the EMDB and the PDB and are publicly available. The accession numbers for maps and models are as follows: the α5β1:BIIG2 complex EMD-44386 and PDB ID 9B9J; the BIIG2 Fab crystal structure: PDB ID 8R38; the α5β1:MINT1526A complex EMD-44387 and PDB ID 9B9K. Two cryoEM data sets have been deposited in the EMPIAR database, each containing raw movies, aligned micrographs and particle stacks. The accession numbers are: α5β1:BIIG2 (EMPIAR-12552); α5β1:MINT1526A (EMPIAR-12553). Analysis scripts for molecular dynamics simulations are available at https://github.com/CampbellLab2020/nguyen_2023. Simulation trajectories and topology files are available on Zenodo at https://doi.org/10.5281/zenodo.10070033. This paper does not report original code. Any additional information required to reanalyze the data reported in this paper is available from the lead contact upon request.

## Acknowledgments

Funding for this research was supported by the National Institutes of Health under Grant K08 CA215105 (A.M.) and R35 GM147414 (M.G.C.). Additional support to M.G.C. was provided by the Seattle Cancer Consortium Safeway Pilot Awards (funded through the NIH/NCI Cancer Center Support Grant P30 CA015704), a Pew Biomedical Scholars award, and Mike and Debbie Koss. A.N. is supported by NHLBI F31 Fellowship 1F31HL174166. MCC was supported by the Interdisciplinary Training Grant in Cancer Research Program (NCI T32 CA080416) through Fred Hutchinson Cancer Center and the Mahan Fellowship; this fellowship is made possible by funding from Mark and Nikki Mahan. J.B.H. was supported by a Mayo Clinic Smith Gibson Research Fellowship award, J.H. by UiO. Part of the work at UiO was funded by Familien Blix fond (grant to H.J. and U.K.). The ALM Holding Company and the Mathy family provided benefactor support.

We thank Genentech and Helen Bettencourt for kindly providing us with purified MINT1526A antibody (proposal OR-217280). We thank the departments and core facilities that supported the generation and analysis of data. Electron microscopy data were generated using the Fred Hutchinson Cancer Center Electron Microscopy shared resource. The EMSR is supported in part by the Cancer Center Support Grant NIH P30 CA15704. We would like to thank Theo Humphreys, Dr. Anvesh Dasari, and Steve MacFarlane for their microscopy assistance and knowledge. We are thankful for the Mayo Clinic Proteomics Core for purification of the Fab components, which is a shared resource of the Mayo Clinic Cancer Center (NIH P30 CA15083). We thank the Fred Hutchinson Cancer Center’s High Performance Computing service for their support in data analysis (1S10OD028685-01). We acknowledge MAX IV Laboratory for time on Beamline BioMax under Proposal 20190447. Research conducted at MAX IV, a Swedish national user facility, is supported by the Swedish Research council under contract 2018-07152, the Swedish Governmental Agency for Innovation Systems under contract 2018-04969, and Formas under contract 2019-02496. At UiO, work was performed at the UiO Structural Biology Core Facilities, which are part of the Norwegian Infrastructure NORCRYST, supported by the Norwegian Research Council (grant 245828). We are grateful to Dr. Thamiya Vasanthakumar, Rachel Werther, Jeremy Hollis, and Andres Fernandez for comments on the manuscript.

## Author Contributions

AN, JBH, MCC, AM and MGC designed the research. CC and ML expressed and purified BIIG2 antibody and prepared BIIG2 and MINT1526A Fab fragments. CC and BM prepared, performed and analyzed mass spectrometry data. AN analyzed the spheroid and tube formation assays. AN, JBH and AM analyzed the tumor xenograft data. JBH performed and analyzed the quantitative PCR experiments. JBH and AM performed and analyzed the immunohistochemistry data. JBH, GC, HJ and UK prepared, collected and analyzed x-ray crystallography data. AN designed, cloned, expressed and purified recombinant α5β1. AN performed and analyzed BLI assays. AN and MGC prepared samples, collected and processed negative stain EM data. AN, CMA and MGC prepared samples and collected cryoEM data. AN and MGC processed the cryoEM data. AN built the cryoEM models. MCC designed, performed and analyzed the molecular dynamics simulations. AN, JBH, AM and MGC wrote the original draft of the manuscript. All authors contributed to reviewing and editing the manuscript.

## STAR Methods

### Key resources table

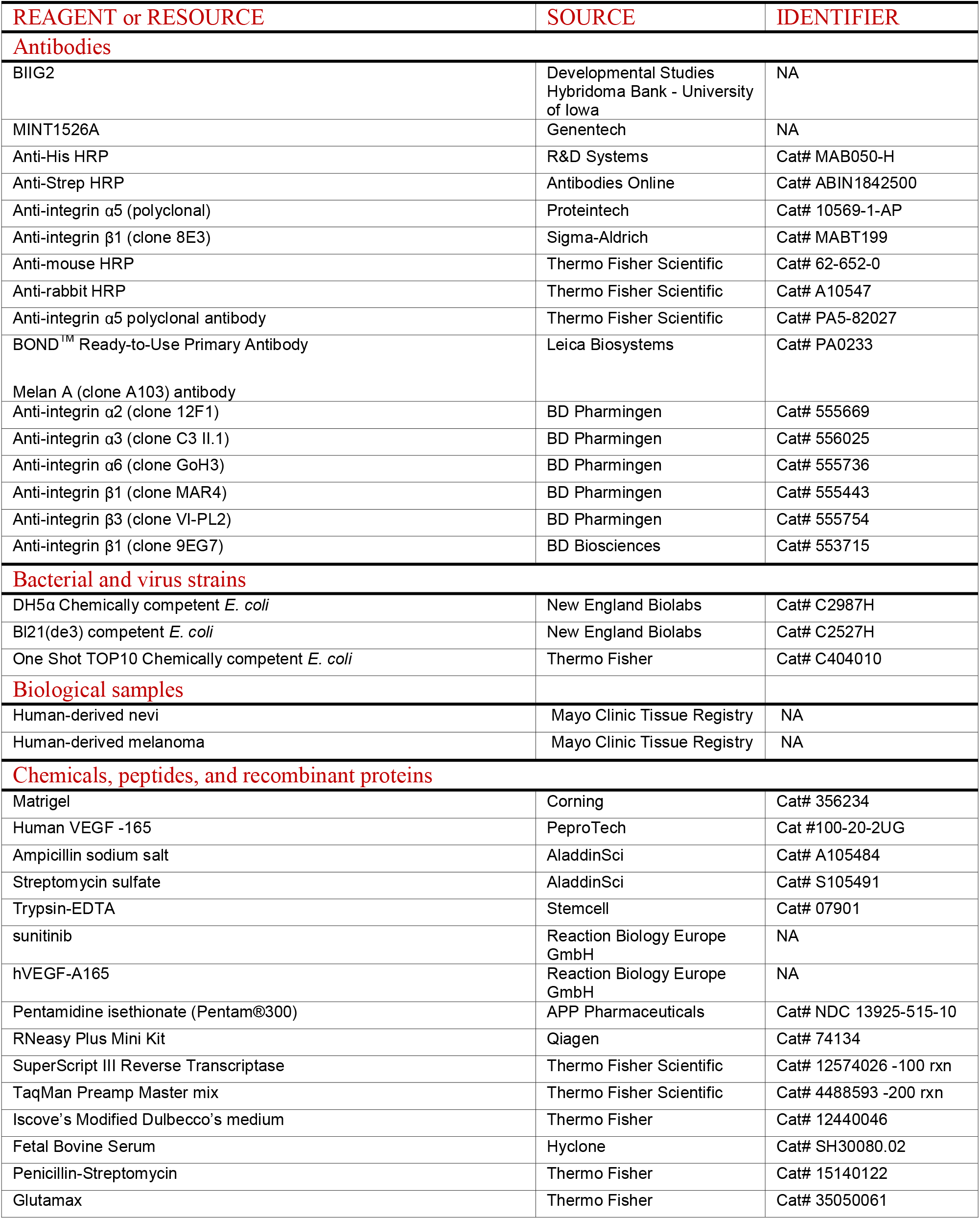
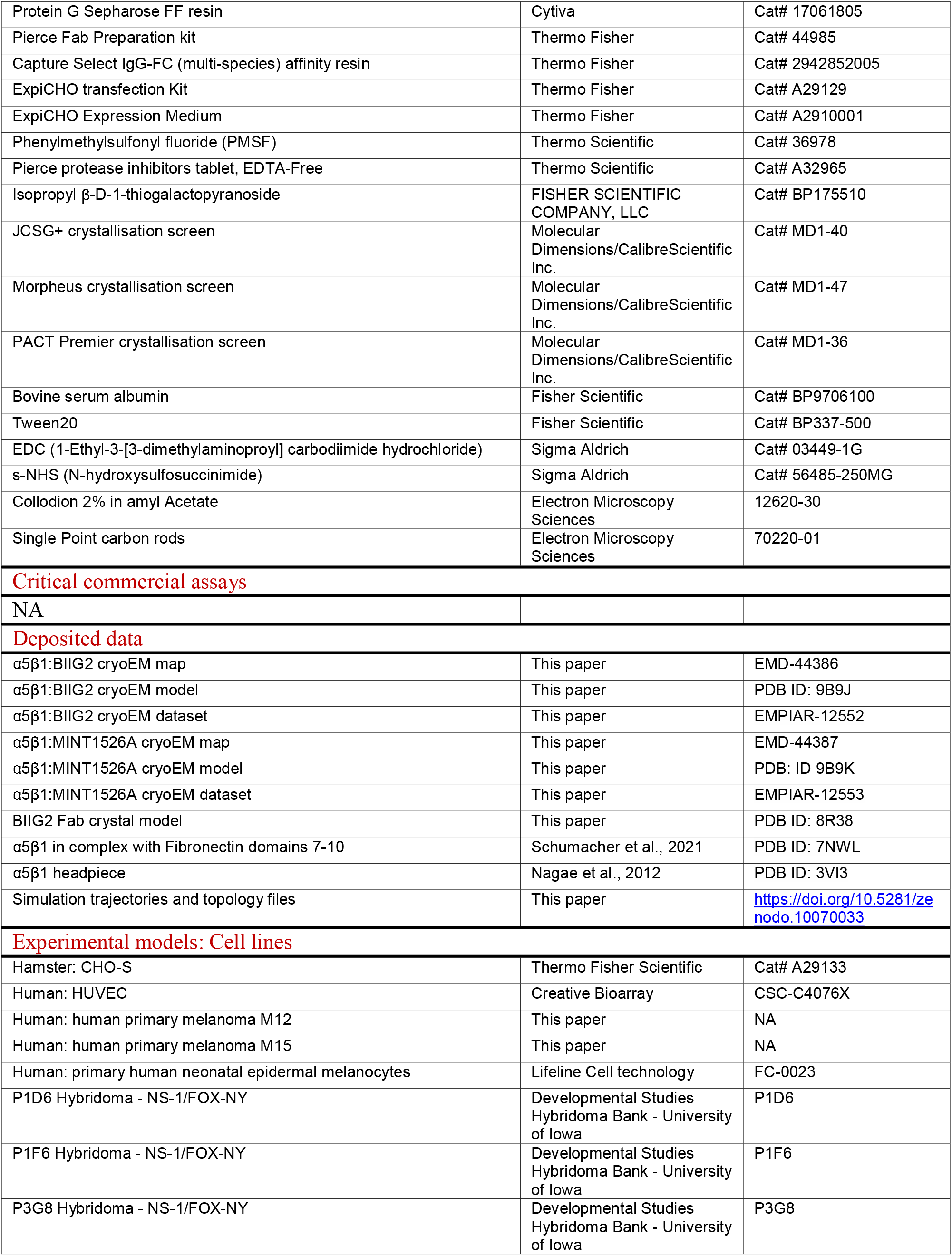
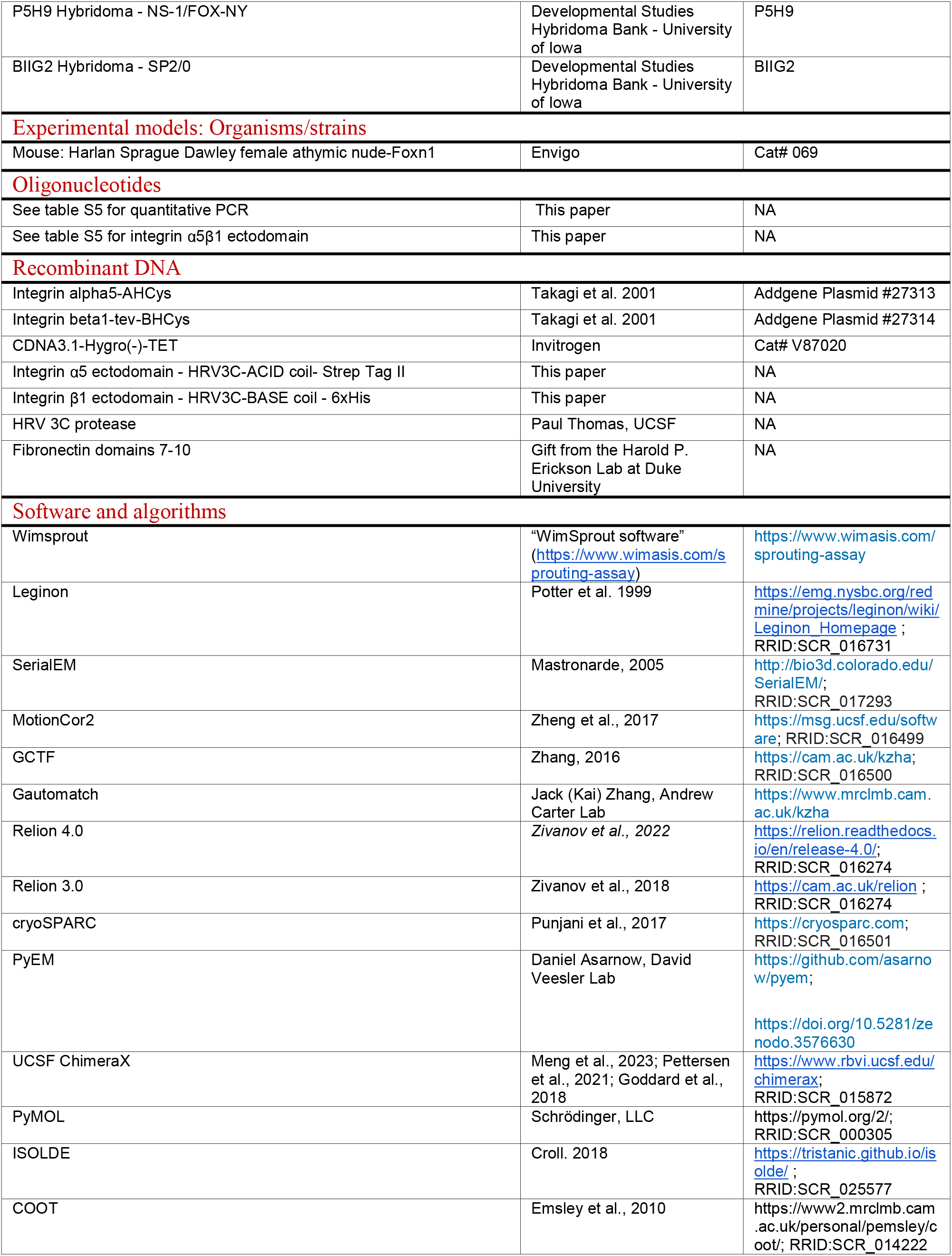
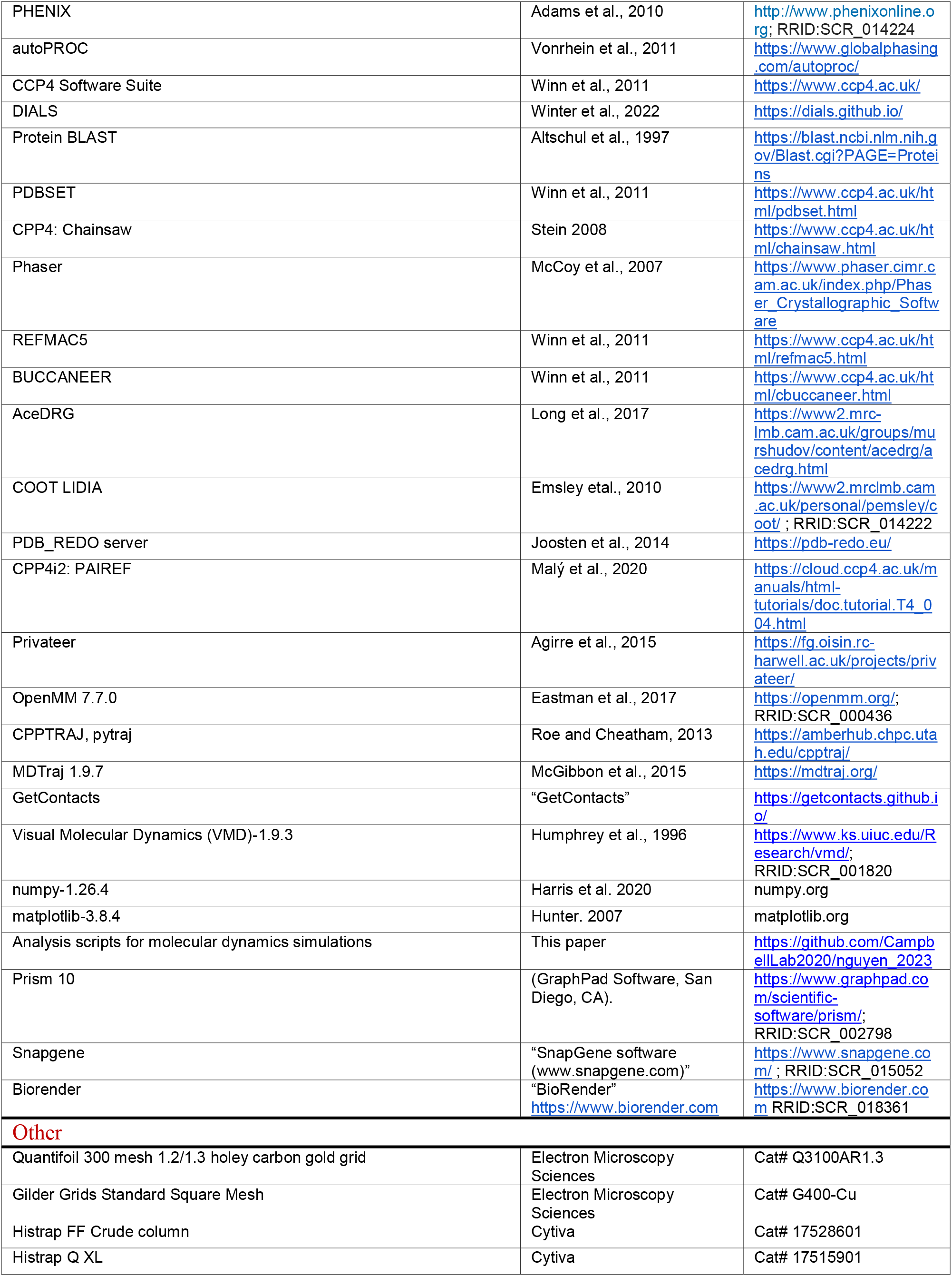
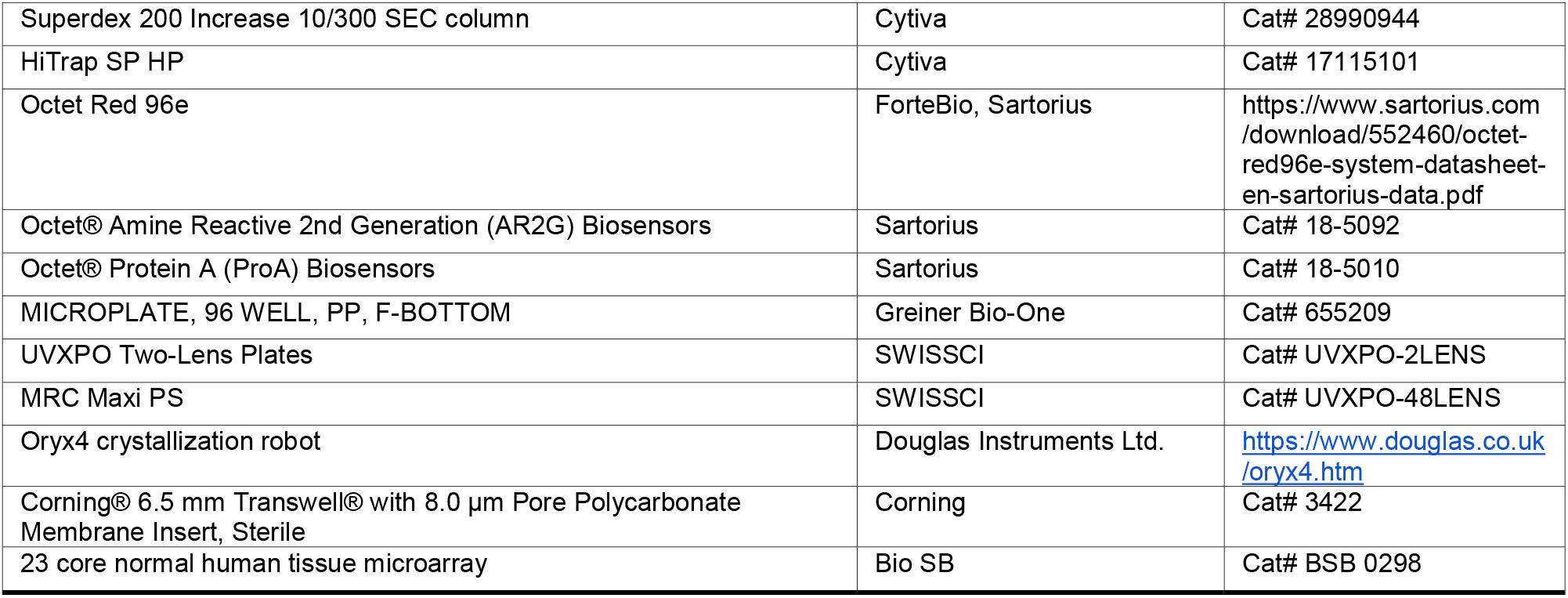

## EXPERIMENTAL MODEL AND STUDY PARTICIPANT DETAILS

### Mouse models

Harlan Sprague Dawley female athymic nude mice that are recessive for the nu mutation in the *Foxn1* gene (Envigo) was used. This mouse strain exhibits a non-functional thymus and are unable to generate cytotoxic effector cells and do not elicit a graft-versus-host response. Mice selected for experimentation were 6 to 7 weeks old, weighing approximately 20 to 30 grams. Their housing conditions involved a 12-hour light/dark cycle with ad libitum access to food and water.

### Cell lines

#### Hybridoma Cell Lines

All hybridomas were purchased from the Developmental Studies Hybridoma Bank. The BIIG2 hybridoma was deposited by Caroline Damsky ^23^. The P1D6, P1F6, P3G8, and P5H9 hybridomas were deposited by Elizabeth Wayner and William Carter ^82,83^. Hybridoma cells were thawed and seeded into 24-well plates with 1 mL of media (IMDM + FBS). Cells were allowed to settle before being placed at 37°C overnight with 5% CO_2_. If cells were too dense, they were further split as described. The following day, cells were pooled from confluent wells and transferred into a T75 flask with 10 mL of media. Cells were passed every other day or when confluent. For antibody expression, cells were seeded onto roller bottles.

#### Patient-Derived Melanoma Cell Lines

Short-term cultures of human primary melanoma cells M12 and M15 were maintained through serial passages in immune-deficient mice via subcutaneous flank implantation, as previously described ^84,85^. Dulbecco’s modified Eagle’s medium supplemented with 10% (v/v) fetal bovine serum (FBS) and antibiotics (penicillin, 100 U/mL, and streptomycin, 100 µg/mL) was used for growing the explant cultures. The cells were kept at 37°C in a humidified incubator with 5% CO_2_.

#### Normal Human Melanocytes

Primary human neonatal epidermal melanocytes were obtained from Lifeline Cell Technology and subsequently cultured using DermaLife M Melanocyte Complete Medium, also provided by Lifeline Cell Technology.

#### Chinese Hamster Ovary Cells

ExpiCHO-S cells were passaged in ExpiCHO Media, cultured at 37°C with 8% CO2 according to the manufacturer’s protocol. Cells were passaged every 3-4 days and were subcultured down to a density of 3.0×10^5^ cells/mL. Before transfection, ExpiCHO-S cells were outgrown to a density of 6.0×10^6^ cells/mL.

### Human tissue samples

A 23 core normal human formalin-fixed paraffin-embedded (FFPE) tissue microarray was purchased from Bio SB (BSB 0298). FFPE tissue of patient-derived nevi and melanoma were obtained through an institutional tissue registry. The Mayo Clinic Institutional Review Board approved the human investigations performed in this study following the Department of Health and Human Services requirements, where appropriate.

## METHOD DETAILS

### 2D-tube angiogenesis formation assay

Endothelial tube formation assays experiments were pursued through Creative Bioarray project CBADY0508. HUVECs (human umbilical vein endothelial cells) were seeded at 5×10^5^ cells/well in 2 mL of complete culture medium in 6-well plates and cultured in a 37°C and 5% CO_2_ incubator. Prior to experimentation, the culture medium was removed. Cells were subsequently treated with 2 mL of culture medium (negative control), 15 ng/mL VEGF (positive control), and BIIG2 or MINT1526A (1, 10, and 100 nM) supplemented with VEGF (15 ng/mL), then incubated at 37°C for 24 hours. Each condition was performed in triplicate. HUVECs were harvested and suspended in serum-free medium and then seeded in Matrigel-coated wells at a density of 1.5×10^5^/1 mL/well. The cells were incubated at 37°C for 6 hours to allow tube formation.

### Spheroid sprouting assay

Experiments were pursued through Reaction Biology under project PQ20385, with modifications of a previously published protocol ^86^. HUVECs were seeded at 1 x 10^4^ cells/cm^2^. HUVEC’s were subcultured every 3-5 days by splitting 1 volume of cells to 3 volumes of ECGM/ECBM media (endothelial cell growth and basal medium, PromoCell) at 37°C with 5% CO_2_. Experiments and spheroid preparation are adapted from previous descriptions ^86,87^. 400 HUVEC were pipetted in a hanging drop to allow overnight spheroid aggregation. 50 HUVEC spheroids were then seeded in 0.9 mL collagen gel and pipetted into a 24 well plate to allow polymerization. The inhibitors combined with 25 ng/mL of VEGF-A were added after 30 minutes by pipetting 100 μL of a 10-fold concentrated working solution on top of the polymerized gel. Plates were incubated at 37°C for 24 hours before being fixed with 4% paraformaldehyde PFA. Sprouting intensity of HUVEC spheroids were captured using an inverted microscope and the digital imaging software NIS-Elements BR 3.0 (Nikon) by determining the cumulative sprout length per spheroid (CSL). 10 randomly selected spheroids were measured and treated as an individual data point.

### Mouse tumor experiment

The mice were randomized into treatment groups and monitored daily. Tumor cells (M12 or M15) embedded in Matrigel were injected subcutaneously into the flank of the athymic mice. In efficacy studies, groups of mice (10 per group) with flank tumors were administered treatment once the tumor volumes became palpable (approximately 50 to 100 mm^3^). The flank tumors were measured three times a week. Treatment was administered twice weekly at a dose of 10 mg/kg. The experiments were concluded when the average tumor volume in the control group exceeded 1,000 mm^3^. All experimental procedures adhered to the Principles of Laboratory Animal Care (NIH) and were approved by the Institutional Animal Care and Use Committee (Mayo Clinic).

### Immunohistochemistry

Proteins of interest were detected using the Leica Biosystems BOND-MAX autostainer platform in combination with primary antibody **(complete list in Key Reagents Table)**, post-primary antibody, and a BOND Polymer Refine Red Detection system (DS9390; Leica Biosystems). Heat-induced antigen retrieval was by BOND EDTA-based pH 9.0 epitope retrieval solution 2 (AR9640; Leica Biosystems) applied for 20 minutes. All stains were digitized with a Leica Aperio ScanScope system.

### Flow cytometry

Normal human melanocytes, M12 and M15 melanoma cells were stained using primary antibodies **(complete list in Key Reagents Table)** against cell surface proteins as shown and PE-labeled secondary antibodies. The BD Bioscience FACScaliber flow cytometer was utilized for data acquisition, and the acquired data was analyzed using FlowJo software.

### Quantitative Microfluidic PCR

Our approach to PCR was as previously described ^88^. RNA from cells was isolated using the RNeasy Plus Mini Kit (Qiagen). One microgram of total RNA was transcribed into cDNA using the SuperScript III Reverse Transcriptase (Thermo Fisher Scientific) and preamplified using TaqMan Preamp Master Mix (Thermo Fisher Scientific). Quantitative reverse transcriptase PCR was then performed using the Fluidigm BioMark HD system and dynamic array-integrated fluid circuits (Fluidigm). After thermal cycling, raw Ct (threshold cycles) data were checked for linear amplification. Gene expression was corrected by the mean of housekeeping genes (RLP0, RLP8, and β-actin) using the ΔCt method. Gene expression input for data analysis was ΔCt. The complete list of primers (HPLC purified, Integrated DNA Technologies) used for cDNA amplification by quantitative PCR are described in **Table S5**.

### Recombinant **α**5**β**1 plasmid design

The sequence of the α5 and β1 subunit wild-type ectodomain was incorporated into a CDNA3.1-Hygro(-)-TET template (Invitrogen). The integrin amino acid numbering starts after the signal peptide. Each integrin subunit was fused with either an ACID or a BASE coiled-coil peptide to assist in heterodimer production ^89^. Residues 1 to 954 of the α5 subunit were cloned followed by a HRV-3C protease cleavage recognition site, an ACID coil, and a Strep-Tag II. The residues 1-728 of the β1 subunit were put into the same plasmid backbone filled by a HRV-3C protease cleavage recognition site, a BASE coil, and a 6x histidine tag. The list of primers used for construct generation are described in **Table S5**.

### Purification of integrin **α**5**β**1 ectodomain

Soluble integrin ectodomain (α5 residues 1-954 and β1 residues 1-728) was expressed using ExpiCHO-S cells (Thermo Fisher) in suspension at a density of 6.0 x 10^6^ cells/mL by transient transfection using the ExpiCHO transfection kit. Cultures were co-transfected with plasmids encoding the α5 or β1 subunit in a 2:1 (α:β) ratio with Expifectamine CHO. Expression was according to the manufacturer’s ‘max titer’ protocol. Cells were harvested by centrifugation at 1,000 x g for 15 minutes at 4°C. Phenylmethylsulfonyl fluoride (PMSF) was added to the supernatant at a final concentration of 0.5 mM before undergoing another centrifugation run at 36,000 x g for 45 minutes at 4°C. Clarified supernatant was diluted 1:1 v/v with Histrap binding buffer (20 mM Na_2_PO_4_, 500 mM NaCl, and 20 mM imidazole, pH 7.4) and applied to 5 mL Histrap FF Crude column (Cytiva) equilibrated with Histrap binding buffer. The column was washed with 1 column volume (CV) of Histrap binding buffer before eluting protein with 2 CV of Histrap Elution Buffer (20 mM Na_2_PO_4_, 500 mM NaCl, and 500 mM imidazole, pH 7.4). Protein fractions were pooled, concentrated and buffer exchanged into integrin storage buffer (20 mM Tris-Base pH 7.4, 150 mM NaCl, 1 mM CaCl_2_, and 1 mM MgCl_2_). Protease was added 1:20 w/w (protease:α5β1) to cleave the ACID/BASE coils and purification tags at 4°C overnight with rocking. The cleaved α5β1 integrin was further purified via gel filtration chromatography using a Superdex 200 Increase 10/300 SEC column (Cytiva) equilibrated in integrin storage buffer. Peak fractions of α5β1 were pooled, concentrated, flash-frozen with 10% glycerol (v/v), and stored at-80°C until used for experiments.

### Purification of BIIG2 and MINT1526A Fab fragments

BIIG2 antibody was produced at scale through the Mayo Clinic institutional core facility. Briefly, the production process involved culturing the hybridoma cells in roller bottles containing Iscove’s Modified Dulbecco’s Medium supplemented with gamma globulin-free serum. A single-step Protein G affinity purification was used to isolate antibodies. The eluted antibody fractions were pooled, concentrated, and buffer exchanged into phosphate buffered saline (1x PBS). The concentrated stock solutions of the antibodies were stored at-80 □C until used for experiments. Genentech has kindly provided us with purified MINT1526A antibody.

BIIG2 and MINT1526A antibodies were digested with the Pierce Fab Preparation Kit (Thermo Fisher) using papain and following the manufacturer’s protocol. BIIG2 Fab fragments were separated from the Fc fragments and intact IgG by loading onto a chromatography column packed with 5 ml of CaptureSelect IgG-Fc (Multi-species) Affinity Resin (Thermo Fisher) in PBS using a NGC FPLC instrument (Bio-Rad). The Fab fragment was present in the flow-through. Separation of the MINT1526A Fab fragment from the Fc fragment was achieved using a NAb Protein A spin column following the manufacturer’s protocol. SDS-PAGE gel bands of Fab and Fc samples from both antibodies were digested and analyzed by mass spectrometry (Mayo Clinic Proteomics Core) to verify the identity and separation of the two fragments.

### Purification of Fibronectin domains 7-10

Human fibronectin domains 7-10 (FN7-10) was purified as described previously ^15^. FN7-10 was expressed in *Escherichia coli* BL21 (DE3) in LB media containing ampicillin at 37 □C with shaking. Cells were grown until the OD600 reached 0.6-0.8. The cells were induced using IPTG at 1 mM final concentration and then grown overnight at 18 □C. Cells were pelleted and resuspended in a buffer containing 50 mM Tris-base pH 7.4, 200 mM NaCl, 1% Triton X-100, and a Pierce protease tablet. Cells were then lysed via sonication and clarified by centrifugation at 76,000 x g for 40 minutes at 4°C. Solid (NH_4_)_2_SO_4_ was added to 40% (final w/v) slowly and the solution was placed on a nutator for 30 minutes at 4°C. This solution was then centrifuged again at 76,000 x G for 40 minutes at 4°C and the pellet was resuspended in Hitrap Q Binding Buffer (20 mM Tris-base pH 7.4 and 50 mM NaCl). The protein sample was applied onto a 5 mL Hitrap Q XL column. The column was washed with 5 CV of Hitrap Q Binding Buffer. The protein was eluted in a gradient profile from 50 mM NaCl to 500 mM NaCl over 20 CV. Protein fractions were pooled, concentrated and buffer exchanged into 50 mM Tris-base pH 7.4 and 150 mM NaCl. FN7-10 was further purified via gel filtration chromatography using a Superdex 200 increase 10/300 SEC column (Cytiva) equilibrated in 50 mM Tris-base pH 7.4 and 150 mM NaCl. Peak fractions of FN7-10 were pooled, concentrated, flash frozen with 10% glycerol (v/v), and stored at-80°C until used for experiments.

### Biolayer Interferometry

All binding experiments were performed on an Octet Red96e (Sartorius). MINT1526A was immobilized on Protein A biosensors (Sartorius). These sensors were hydrated using nonactive running buffer (20 mM Tris-Base pH 7.4, 150 mM NaCl, 0.1% BSA, and 0.02% Tween20 with 5mM CaCl_2_) for 15 minutes before subsequently being incubated with fresh nonactive running buffer. MINT1526A was diluted with nonactive running buffer to 5 ug/mL and loaded for 300 seconds before a 120 second baseline equilibration in the nonactive running buffer. Association with integrin α5β1 ectodomain was carried out with a two-fold dilution series from 50nM to 1.56nM in nonactive running buffer for 300 seconds before dissociating for 300 seconds. Experiments were repeated with active running buffer (20 mM Tris-Base pH 7.4, 150 mM NaCl, 0.1% BSA, and 0.02% Tween20 with 1mM MnCl_2_). MINT1526A binding experiments were performed in triplicate for both nonactive and active conditions.

Binding assays with BIIG2 began with hydrating AR2G biosensors (Sartorius) in water for 15 minutes, followed by an additional 1 minute incubation. The AR2G biosensors were then activated using 20 mM EDC and 10 mM s-NHS for 300 seconds (following manufacturer’s protocol). BIIG2 antibody was diluted with 10 mM sodium acetate pH 5.0 to 5 ug/mL and loaded until a threshold shift of 2nm was achieved. Loading was immediately quenched in 1M ethanolamine for 300 seconds. A baseline equilibration step was performed in nonactive running buffer for 120 seconds. Association with integrin α5β1 ectodomain was performed at a two-fold dilution series from 50nM to 1.56nM in nonactive running buffer for 450 seconds before dissociating for 450 seconds. Experiments were repeated with active running buffer and each buffer condition was repeated for an additional two times.

Competitive binding assays were carried out where α5β1 was bound to MINT1526A or BIIG2 Fabs or FN domains 7-10 fragment in a 1:2 molar ratio 20 minutes prior to assay running in either active or nonactive buffer. MINT1526A or BIIG2 antibodies or FN 7-10 used as immobilized ligands were loaded onto their respective biosensors (Protein A for MINT1526A and AR2G for BIIG2 and FN 7-10) at 5ug/mL. Experimental conditions and association and dissociation step times follow as described above.

### Crystallization of BIIG2

BIIG2 Fab (7.14 mg/mL in PBS + 0.02% sodium azide) was diluted in CIEX binding buffer (25□mM Bis-Tris-HCl, pH 6.5) and loaded onto a pre-equilibrated 1 mL HiTrap SP HP chromatography column (Cytiva), connected to an ÄKTA Purifier 10 chromatography system (Cytiva). The protein was eluted with a 0-100% gradient over 100 mL of CIEX elution buffer (25□mM Bis-Tris-HCl, 1□M NaCl, pH 6.5). The main protein component eluted as a highly symmetric peak around 12% v/v Buffer B (approximately 120 mM NaCl). Fractions containing BIIG2 Fab protein were pooled and concentrated to 10 mg/mL for crystallization studies.

Preliminary screening of BIIG2 Fab crystallization conditions was performed with pre-formulated commercial screens (JCSG+, Morpheus, PACT Premier, Molecular Dimensions/CalibreScientific Inc.) ^90–92^. Sitting-drop vapor diffusion crystallization experiments were set up on UVXPO Two-Lens Plates (SwissCI) by mixing BIIG2 Fab at 10 mg/mL and the crystallization solutions in a 1:1 volume ratio (300 nL + 300 nL) using an Oryx4 crystallization robot (Douglas Instruments Ltd.) Plates were stored at 20°C in a RI-182 crystal plate hotel (Formulatrix), and monitored at regular time intervals for the formation of crystals. Initial crystals grew from several conditions from the JCSG+, PACT Premier and Morpheus formulations. Two sets of 48 optimized crystallization solutions were designed, combining the chemical components from several PACT Premier hits. A new sitting-drop vapor diffusion experiment was set up by mixing the optimized formulations with CIEX-purified BIIG2 Fab, which was diluted with type I water (Milli-Q) to a protein concentration of 7.6 mg/mL.

Crystallization drops were set up on an MRC Maxi PS (SwissCI) crystallization plate, mixing the diluted protein solution and the crystallization cocktails in a 1:1 v/v ratio (300 nL + 300 nL). Diffraction-quality crystals grew from several conditions. Crystals with the most promising morphologies were harvested in nylon loops or lytholoops, cryoprotected (if appropriate), flash-cooled in a nitrogen cryostream, and sent for synchrotron data collection in a liquid nitrogen-cooled CX 100 dry shipper (Taylor Wharton). The crystal that provided the highest-diffracting data set grew from 20% w/v PEG 3350, 0.2 M sodium chloride, 2.5% v/v DMSO; cryoprotectant solution was prepared by complementing the crystallization solution from the reservoir with glycerol, to a final concentration of 20% v/v.

### X-ray data collection and refinement

Synchrotron data collection on BIIG2 Fab crystals was performed at the BioMAX beamline, MAX IV, Lund, Sweden (at 100□K, 0.9537□Å). Data were initially integrated and scaled with *autoPROC* ^93^, and ranked by their diffraction resolution cut-off, based on the assessment of the Pearson’s correlation coefficient (CC_1/2_ > 10%) ^94^.The highest-resolution data set (1.4□Å), collected on a crystal belonging to space group *P*2_1_2_1_2_1_, was chosen for structure determination. Many of the programs and tools used belong to the *CCP4* software suite ^95^. Data were integrated and scaled with *DIALS* ^96^; merging and truncation were carried out with *AIMLESS*, a component of the *CCP*4 software suite ^95^. A PDB-restricted BLAST ^97,98^ homology search identified a suitable search model for molecular replacement, the Fab fragment of a rat monoclonal antibody (PDB ID: 1C5D; VL=80.5 % seq. id.; VH = 79.1% seq. id.). Ligands and water molecules were manually deleted, and alternative conformations removed using PDBSET ^95^. Amino acids were truncated to the last conserved atom using the *CCP*4 tool *CHAINSAW* ^99^. The truncated model was segmented into four separate components, the heavy and light chains and their constant and variable regions, which were used as separate search models in *Phaser* ^100^. The phased data set contained a single copy of the Fab heterodimer in the asymmetric unit. Search model bias was mitigated by running an extensive initial round (60 cycles) of maximum-likelihood refinement using *REFMAC*5 ^95^. Two cycles of real-space model rebuilding were carried out in *Coot* ^101^, each followed by maximum-likelihood isotropic refinement with *REFMAC*5 ^95^. Missing atoms from the protein main chain were automatically added with *BUCCANEER* ^95^, followed by several rounds of manual building with *Coot* ^101^, alternating with maximum-likelihood isotropic Single-wavelength anomalous diffraction (SAD) refinement with *REFMAC*5 ^95^. The final refinement step included TLS parameterization ^102^.

After correcting inaccuracies in the main protein chain, first and second-coordination sphere water molecules were added to the model, followed by alternative main and side-chain conformations. Water molecules were initially placed using *Coot*:Find waters and then manually inspected for several criteria, including distances from hydrogen-bond donors/acceptors and quality of the electron density. Occupancies for different side chain conformations were assigned during the course of a combined real-space/maximum-likelihood refinement round through *phenix.refine* ^103,104^. The model was completed by adding ligands, including part of the *N*-linked glycosyl chain visible on N60 and glycerol, which had been used as cryoprotectant. Well-defined saccharide units from the glycan core region were manually positioned in the difference density map and linked by editing the coordinates file. Glutamine H1, the N-terminal amino acid of the Fab heavy chain, was modeled as pyroglutamate. Some of the disulfide bridges were found to be partly reduced (*e.g.*, C23-C88, chain L), due to minor radiation damage; the relevant cysteine residues were modeled as partly reduced, when appropriate. K199 showed residual density on the terminal amino group of its side chain and was modeled as methyllysine. His H100C showed residual density suggesting post-translational modification, and was modeled as 5-hydroxyproline. As this residue is not included in the *REFMAC* dictionary, a ligand definition file was created with *AceDRG* ^105^. 5-hydroxyproline (PubChem CID: 14325405) coordinates were generated with *Coot LIDIA* ^106^, starting from an isomeric SMILES string obtained from the PubChem server ^107^. The *AceDRG* output for the new ligand (HY5) was edited, renaming and renumbering atoms to match the standard proline numbering; occupancy refinement suggested an 86% post-translational modification rate for His H100C. Further model improvements were assessed with the PDB_REDO server ^108^, while the diffraction limit was validated by running paired refinement using *PAIREF* ^109^, accessed through the *CCP4i2* GUI ^110^. The overall quality of the model was validated through the PDB Validation server ^111^. The percentages of amino acid residues occupying the favored, allowed and outlier regions in the Ramachandran plot are 98/2/0%. The quality of the carbohydrate chain was evaluated by *Privateer* ^112^, a *CCP4* tool accessible through the *CCP4i2* interface ^110^. σ_A_-weighted electron density maps were generated using the FFT tool, a component of the *CCP4* software package ^95^. Figures were generated using PyMOL (Schrödinger Inc.), assigning secondary structure elements as defined by the SST web server ^113^. Position numbering in the final model was adjusted to match the Kabat scheme ^114^, before depositing the refined coordinates and structure factors in the PDB ^115^, with PDB ID: 8R38.

### Negative-stain EM sample preparation, data acquisition and processing

The α5β1-Fab complexes were formed using a 1:2 molar ratio of α5β1 to antibody Fab, incubated at room temperature for 20 minutes, and diluted to a final concentration of 7.5 ug/mL in 20 mM Tris-base pH 7.4, 150 mM NaCl supplemented with either 5 mM CaCl_2_ or 1 mM MnCl_2_. For experiments determining what integrin conformations MINT1526A binds, we used the commercially available conformation-inducing antibody, 9EG7 IgG (BD Biosciences cat# 553715). Integrin α5β1 ectodomain was buffer-exchanged into nonactivating conditions (5 mM Ca^2+^, 20 mM Tris-base pH 7.4, 150 mM NaCl) for 10 minutes with rocking at room temperature. 9EG7 IgG was added to integrin α5β1 at a 1:1 molar ratio and incubated for 10 minutes at room temperature with rocking. 9EG7 binding locks α5β1 into the extended state ^43^, then MINT1526A or BIIG2 Fab was added at a 1:1 molar ratio to integrin α5β1 ectodomain and incubated for an additional 10 minutes with rocking at room temperature. Experiments were repeated in activating buffer (1 mM Mn^2+^, 20 mM Tris-base pH 7.4, 150 mM NaCl).

For all experiments, 3 µL of protein sample was added to a glow-discharged (PELCO easiGlow; 30 seconds; 15 mA) carbon-coated copper grid (homemade carbon layer). The specimens were stained with 3 µL of 2% (w/v) uranyl formate solution as previously described ^116^. Data were acquired using a Thermo Fisher Scientific Talos L120C transmission electron microscope operating at 120kV and recorded on a 4k x 4k Thermo Fisher Scientific Ceta camera at a nominal magnification of 92,000x with a pixel size of 1.58 Å/px. Leginon ^117^ was used to collect 252 (apo α5β1 nonactivating), 253 (apo α5β1 activating), 385 (α5β1:BIIG2 nonactivating), 399 (α5β1:BIIG2 activating), 422 (α5β1:MINT1526A nonactivating), 482 (α5β1:MINT1526A activating), 368 (α5β1:9EG7:BIIG2 nonactivating), 373 (α5β1:9EG7:BIIG2 activating), 366 (α5β1:9EG7:MINT1526A nonactivating) and 395 (α5β1:9EG7:MINT1526A activating) micrographs with a nominal range of 1.7-2.0 µm under focus and a dose of approximately 50 e^-^/Å^2^. The contrast transfer function of each micrograph was estimated using GCTF ^118^, and particles were automatically picked unbiasedly using Gautomatch ^118^ (http://www.mrc-lmb.cam.ac.uk/kzhang/Gautomatch). Particles were extracted at 352 pixels and binned to 128 pixels using RELION3 ^119^ before being imported into cryoSPARC ^120^. Particles were subjected to multiple rounds of reference-free 2D alignment and classification to remove false-positive particle images. The final particle count contributed to 2D class averages were 26,057 (apo α5β1, nonactivating), 31,786 (apo α5β1, activating), 18,200 (α5β1:BIIG2, nonactivating), 16,797 (α5β1:BIIG2, activating), 9,742 (α5β1:MINT1526A, nonactivating), 41,387 (α5β1:MINT1526A, activating), 21,329 (α5β1:9EG7:BIIG2 nonactivating), 25502 (α5β1:9EG7:BIIG2 activating), 12,149 (α5β1:9EG7:MINT1526A nonactivating) and 48,765 (α5β1:9EG7:MINT1526A activating).

### Cryo-EM sample preparation

α5β1-BIIG2/MINT1526A Fab complexes were formed using a 1:2 α5β1 to Fab molar ratio in nonactivating buffer (20 mM Tris pH 7.4, 150 mM NaCl, 5 mM CaCl_2_) at room temperature with rocking for 20 minutes. nonactivating conditions were used to drive the equilibrium toward a more homogenous sample since we had already established that the epitope binding site was nowhere near the relevant cation coordination sites. These protein complexes were subjected to size exclusion chromatography using a Superdex200 Increase 10/300 GL (Cytiva) equilibrated with nonactivating buffer. Fractions of α5β1:BIIG2/MINT1526A Fab were pooled and concentrated to 0.15 mg/mL before 3 uL of the protein sample was placed on a glow discharged (PELCO easiGlow; 30 seconds; 15 mA) R1.2/1.3 Au 300-mesh grid (QUANTIFOIL), blotted for 8 seconds with a blot-force of 1, and plunge-frozen immediately in liquid ethane using a Thermo Fisher Scientific Vitrobot Mark IV at 4° C and 100% humidity.

### Cryo-EM Data collection and processing

Two datasets (one each for α5β1:BIIG2 Fab and α5β1:MINT1526A Fab) were acquired on a Thermo Fisher Scientific Glacios cryo-transmission electron microscope operating at 200kV and recorded with a Gatan K3 Direct Detection Camera. Automated data collected was carried out using the SerialEM software ^121^. 99 frame movies were recorded in super-resolution mode with a super-resolution pixel size of 0.561 Å/px at a nominal magnification of 36,000x. Both datasets were collected in a single session with a nominal defocus range of 1.4–1.8 μm under focus and a dose of approximately 50e^-^/Å^2^. 3656 micrographs were collected of the α5β1:BIIG2 Fab complex and 3079 micrographs of α5β1:MINT1526A Fab.

The dose fractionated super-resolution image stacks were motion corrected and binned 2×2 by Fourier cropping using MotionCor2 ^122^. Motion corrected images were then processed using CTFFIND4 within the cryoSPARC wrapper ^120,123^. 2,425,530 (α5β1:BIIG2 Fab) and 1,101,393 (α5β1:MINT1526A Fab) particles were initially picked using the unbiased blob picker in cryoSPARC. These particles were subjected to multiple rounds of reference-free 2D alignment and classification in cryoSPARC. Next, particles were polished in Relion ^124,125^ and final 3D classification and refinement steps were done in cryoSPARC. The final particle count for α5β1:BIIG2 Fab is 371,066. The final particle count for α5β1:MINT1526A Fab is 342,608. Images showing EM maps were generated using UCSF ChimeraX ^126–128^. A detailed processing schematic is in **Figure S10**.

### CryoEM model building

The closed α5β1 headpiece (PDB: 3VI3) and our solved BIIG2 Fab structure were used as initial models for the α5β1:BIIG2 Fab complex. The half bent α5β1 ectodomain (PDB ID: 7NXD) was used as an initial model for α5β1:MINT1526A Fab complex. MINT1526A was modeled *de novo* using peptide sequences derived from mass spectrometry. Initial models were fitted into their respective cryoEM density using UCSF ChimeraX ^126–128^ and manually adjusted in COOT ^101^. Glycosylations were manually added using COOT. Models (including glycans) were refined using Phenix and ISOLDE ^104,129,130^. Modeling was aided by using sharpened and unsharpened cryoEM maps. BIIG2 and MINT1526A were numbered according to the Kabat numbering scheme ^131^. CryoEM maps and model statistics are found in **Table S3**.

### Molecular dynamics simulations

All-atom molecular dynamics simulations were performed using the OpenMM 7.7.0 package ^132^ using AMBER ff19SB parameters and topologies. The initial structure for simulations were constructed from the α5β1:BIIG2 or α5β1:MINT1526A cryoEM models. To reduce the system size, only the headpiece domains of α5β1 and variable domains of BIIG2 and MINT Fabs were simulated. N- and C- terminal residues were capped with acetyl and methyl amide groups, respectively. Protonation states for titratable residues were modeled based on a pH of 7.4. The systems were solvated in a truncated octahedron with the minimal distance between the solute and the box boundary set to 8Å and 150 mM NaCl. Calcium ions resolved in the cryoEM model were retained. The system was parameterized using the AMBER19ffSB ^133^ force field combined with the OPC water model and 12-6-4 Li Merz ion parameters ^134^. As a control, an α5β1 without Fab bound, referred to as apo α5β1, was modeled and parameterized similarly.

The systems were first energy minimized for 20,000 steps followed by a gradual heating to 298K. During minimization and heating, protein backbone atoms were harmonically restrained by a 5 kcal/mol*Å^2^ force constant. Production simulations were performed using periodic boundary conditions, hydrogen mass repartitioning ^135^, and a Langevin integrator with an integration timestep of 4 fs and collision rate of √2 ps^-1^. Pressure was maintained at 1 atm using a MonteCarlo barostat with an update frequency of 100 steps. Nonbonded interactions were calculated with a distance cutoff of 10Å. Trajectory snapshots were saved every 100 ps during production simulations. A total of 7 replicates, 2µs each in length, were simulated for α5β1 systems.

Trajectories were processed using in-house scripts utilizing the CPPTRAJ ^136^, MDTraj 1.9.7 ^137^, GetContacts (https://getcontacts.github.io/), numpy-1.67.4 ^138^ and matplotlib-3.8.4 packages ^139^. Hydrogen bond frequencies were calculated using a 3.5 Å distance cutoff between the donor-hydrogen and acceptor atoms. The first 50 ns of production simulations were not included in analysis. Trajectories were visualized using VMD 1.9.2 ^140^.

## QUANTIFICATION AND STATISTICAL ANALYSIS

Spheroid assays are reported as means ± SEM. 8-10 spheroids were measured at random per condition and concentration. Cumulative sprout length per spheroid (CSL) was quantified using Wimsprout image analysis software (https://www.wimasis.com/sprouting-assay). Significance was determined by a measurable reduction of total sprouting length compared to the stimulated sprouts (+VEGF), with adjusted P values less than 0.05. Statistical analysis was performed using a one-way ANOVA with Bonferroni correction post-hoc. Statistical analysis can be found in the legend of **Figure S2**.

2D angiogenesis tube formation assays are reported as means ± SEM. The total master segments length (pixels) and the number of meshes/junctions were counted using Angiogenesis Analyzer ImageJ software^81^. Three representative images were taken for each well at 40x magnification under an inverted microscope and the mean measurement was used as the individual data point. This was repeated for a total of three wells. Statistical analysis was done using a one-way ANOVA with a Bonferroni correction against the control group (HUVEC + VEGF). Significance was determined by a measurable reduction of total tube length and mesh count compared to the VEGF-stimulated control, with adjusted P values of less than 0.05. Statistical analysis can be found in the legend of **Figure 1**.

Mouse tumor models are reported as means ± SEM. Total number of mice per antibody per time point equals n = 8-10. Statistical analysis performed was a one-way ANOVA with Bonferroni correction against the control group (IgG) and significance was determined if any antibodies tested are capable of reducing tumor volume compared to the control IgG with an adjusted P value less than 0.05. Additional one-way ANOVA testing of the other α5 and αv targeting antibodies against BIIG2 was performed with a Bonferroni correction. Significance was determined if BIIG2 performed better than the panel of antibodies at reducing tumor volumes with an adjusted P value less than 0.05. In experiments with Pentamidine, the means of all conditions were compared to one another via one-way ANOVA with a Bonferroni correction. Total number of mice per experimental group is n = 8-10. Significance was determined if any of the treatment conditions were able to reduce tumor volume significantly with an adjusted P value less than 0.05. Statistical analysis can be found in the legend of **Figure 2** and **Figure S5**.

X-ray crystallography and cryo-EM data collection and refinement statistics are summarized in Table S2 and Table S3.

All statistical analyses were performed using the software package Prism 10 (GraphPad Software, San Diego, CA).

