## Supplemental Materials for "Shared ligand-blocking mechanism but distinct conformational modulation by α5-targeting antibodies BIIG2 and MINT1526A"

**This PDF file includes:**

Figures S1 to S13

Tables S1 to S5

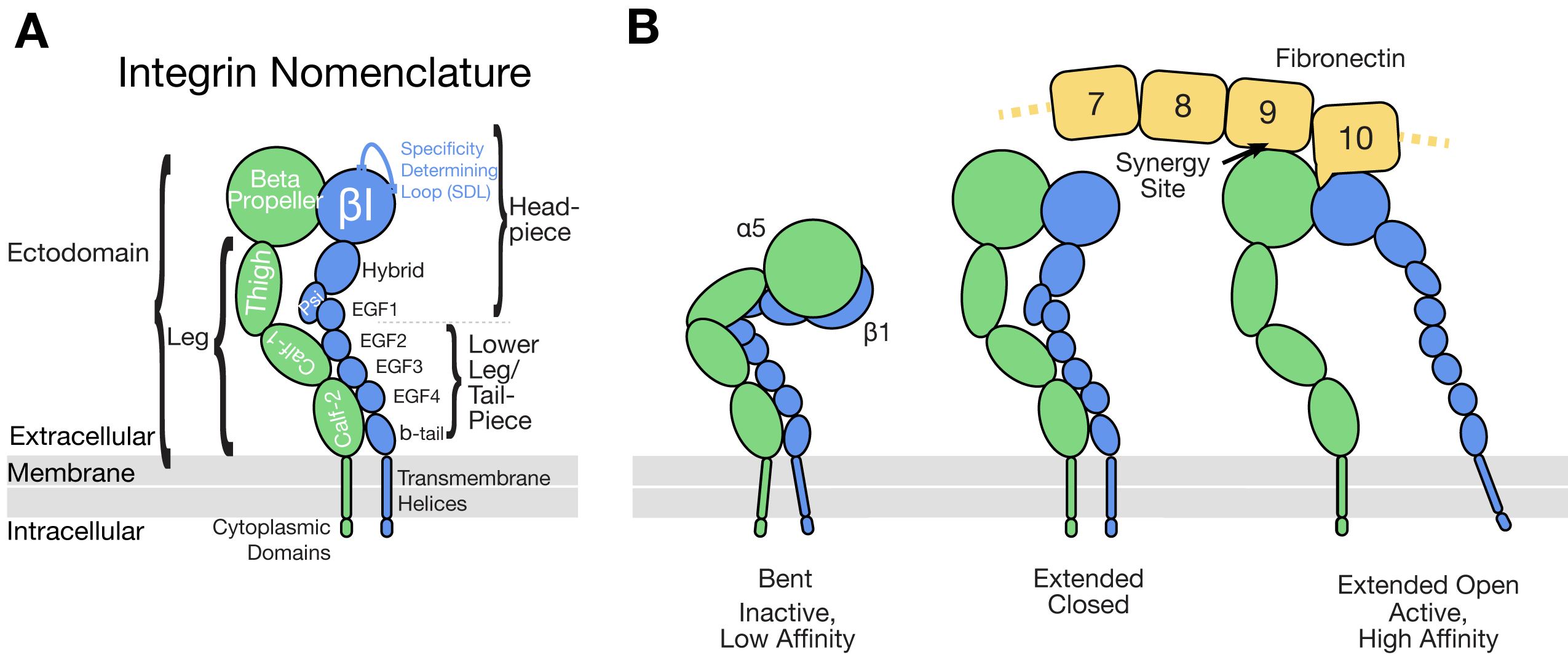

**Figure S1. Integrin and fibronectin nomenclature and conformations, Related to Figure 3. (A)** Integrin designations and domains: The α5 subunit is shown in green, the β1 subunit is shown in blue. **(B)** Integrin α5β1 conformations: bent (half-bent), extended closed, and extended open. Extended open integrin is shown bound to a fragment of fibronectin (domains 7-10, referred to as FN7-10, yellow).

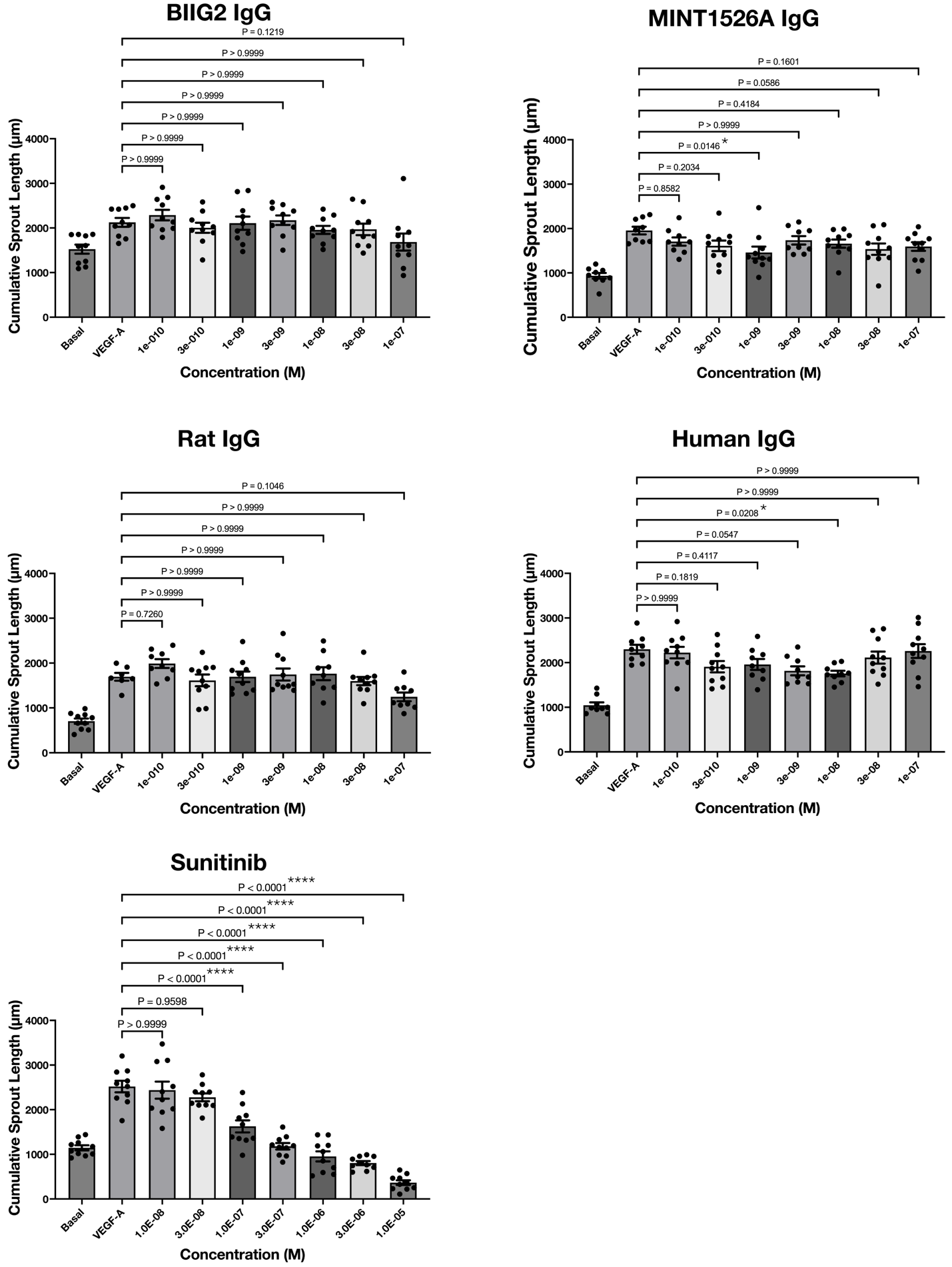

**Figure S2. BIIG2 and MINT1526A does not inhibit spheroid sprouting, Related to STAR Methods.** HUVEC spheroids were imaged and measured for cumulative sprout length (CSL) in μm using WimSprout analysis software. Concentration of inhibitors are in the presence of 25 ng/mL VEGF-A. Basal conditions do not have VEGF-A. Data shown are mean ± SEM, n= 9-10 spheroids. Spheroids in which sprouting could not be assessed are excluded. One-way ANOVA was used to determine whether any concentration of antibody or Sunitinib was capable of reducing VEGF-stimulated spheroid sprouting against the control group (25 ng/mL VEGF-A). Multiple comparison testing was used with Bonferroni’s correction. Adjusted P values are shown, and significance is determined by values less than 0.05.

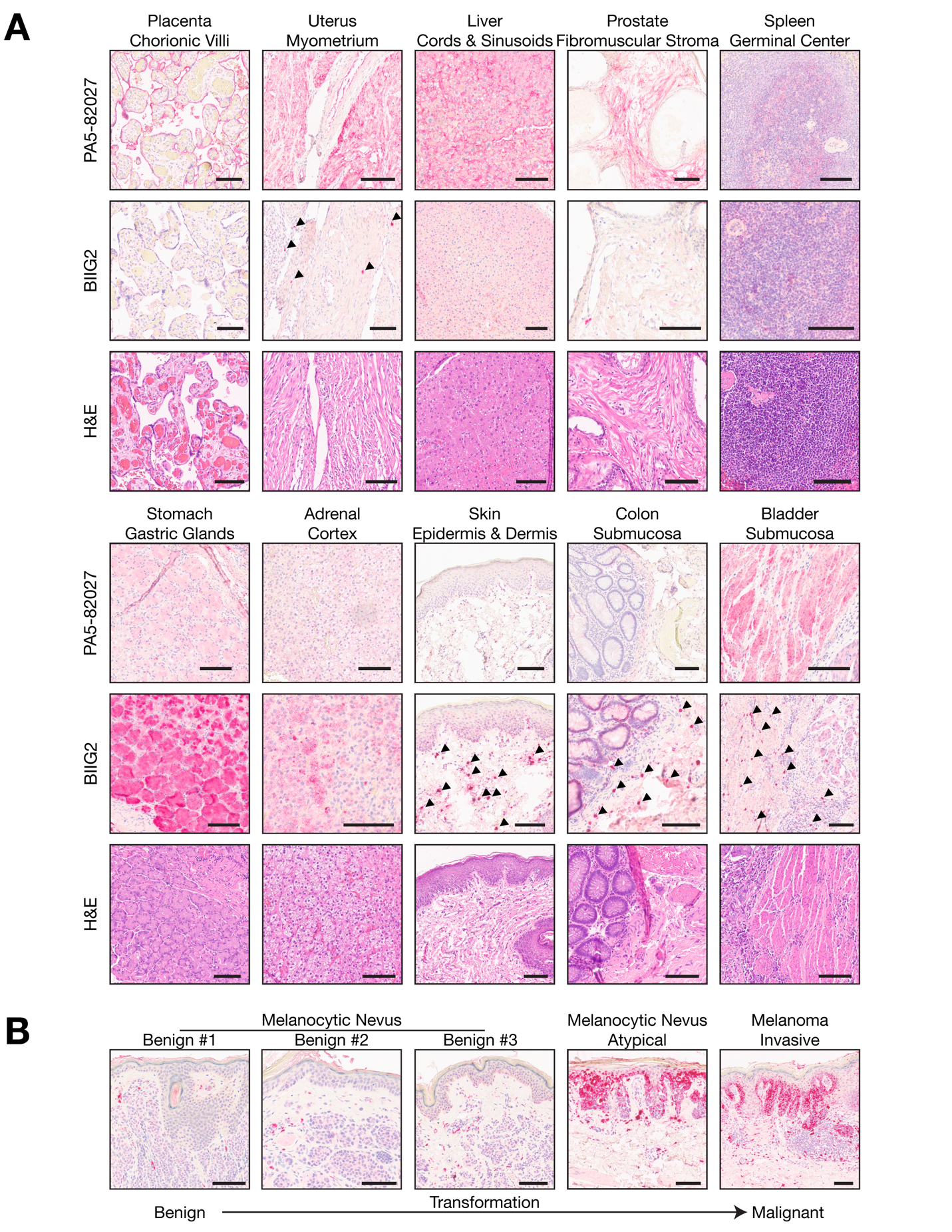

**Figure S3. Staining pattern of BIIG2 and polyclonal α5 integrin antibody PA5-82027 in diverse human tissue samples, Related to STAR Methods. (A)** Immunostaining was performed on consecutive sections from a tissue microarray comprising 23 paraffin-embedded, formalin-fixed (FFPE) core samples of normal human tissues using BIIG2 or PA5-82027 as primary antibodies. Hematoxylin and Eosin (H&E) staining was also performed to enhance the visualization of tissue architecture. Arrows indicate specific clusters of fibroblasts within connective tissues that exhibit intense staining with the BIIG2 antibody, highlighting these clusters as regions of interest. In the zona fasciculata of the adrenal gland, spongiocytes—cells rich in lipid droplets—displayed a speckled staining pattern. Gastric glands, containing proteins and mucus, also showed staining with the BIIG2 antibody, but it appeared non-localized. The speckled and non-localized staining patterns observed in spongiocytes and gastric glands suggest potential non-specific staining. **(B)** Patient-derived biopsies of normal nevi, atypical nevi, and invasive cutaneous melanoma, representing the spectrum of melanocyte malignant transformation, reveal intense BIIG2 staining in atypical melanocytes and melanoma cells within the epidermis and upper dermis. In contrast, benign nevus samples exhibit sparse staining, with only faint staining observed in platelets and vasculature. Scale bars, 100 μm.

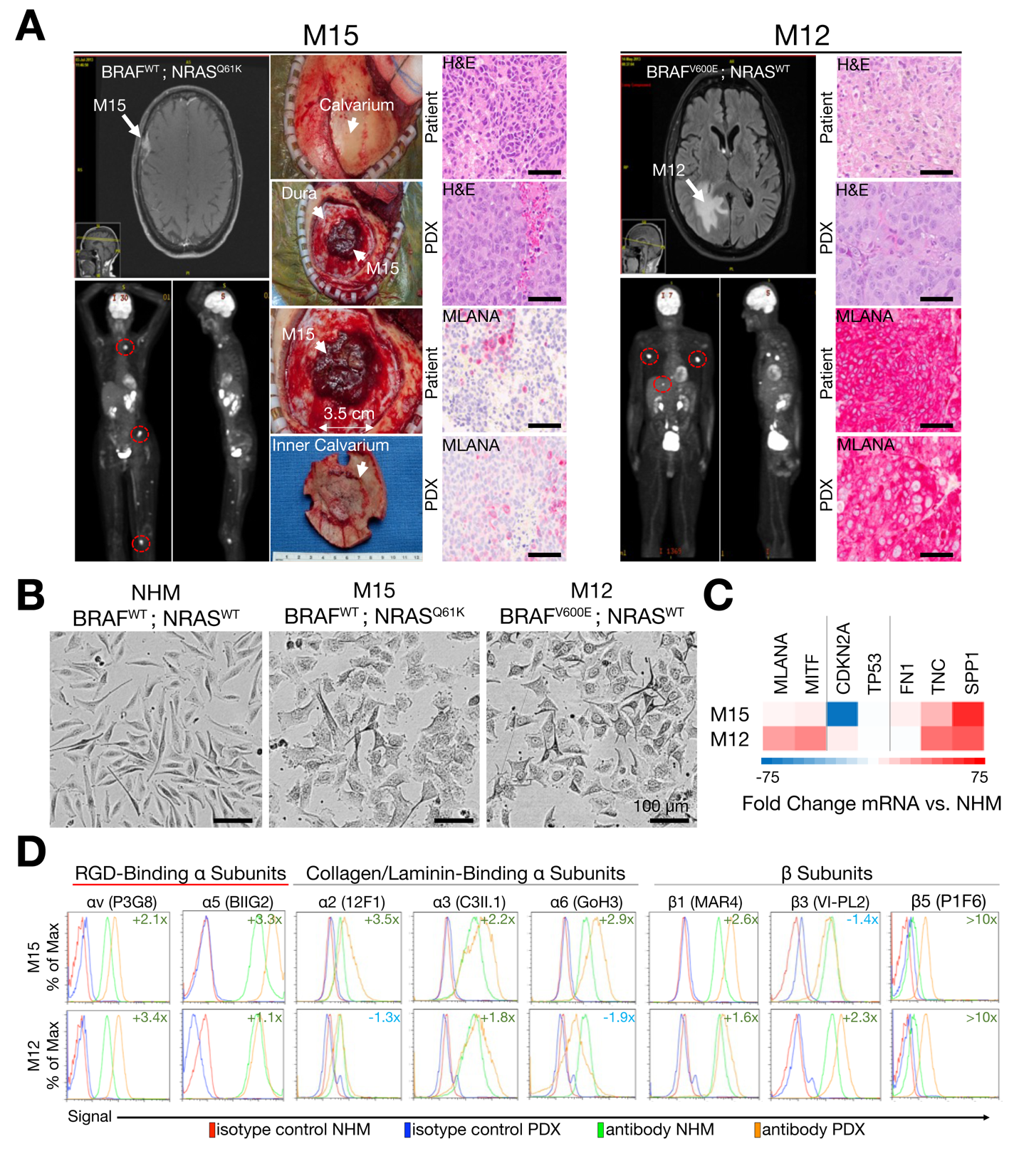

**Figure S4. Characterization of patient-derived xenografts of metastatic melanoma for evaluating the efficacy of integrin antibodies on tumor growth, Related to Figure 2.** **(A)** M15 cultures were established from a 66-year-old female patient with metastatic melanoma, including brain metastases. The M15 xenograft specifically originated from a right dural-based brain mass with involvement of the overlying calvarium (the dome-like upper part of the skull), as identified through magnetic resonance imaging (MRI; indicated by the arrow labeled M15). Positron emission tomography (PET) imaging further revealed diffuse metastases (some areas of metastasis are marked by red circles). Surgical images depict the exposed calvarium, and upon its removal, the melanoma brain metastasis becomes visible, surrounded by the dura mater (the outermost layer of the meninges). Gross examination revealed that the metastasis had a diameter of approximately 3.5 cm. Images from the surgery also show the inner surface of the calvarium. Histological analysis of the patient's brain tumor and the M15 xenograft grown in mice showed similar characteristics in H&E-stained sections, including densely packed nuclei and occasional mitotic figures. Furthermore, protein expression of MLANA (Melan-A), a melanocyte differentiation marker, was similarly sparse in both the patient's brain tumor and the M15 xenograft. This similarity highlights the fidelity of the PDX model in preserving essential tumor characteristics. The patient’s disease began with a primary melanoma on the left ankle, accompanied by microscopic involvement of regional lymph nodes. Over time, the patient developed widespread metastases that did not respond to radiation therapy or immunotherapy with ipilimumab, a monoclonal antibody that targets CTLA-4 (cytotoxic T-lymphocyte-associated protein 4). The patient passed away two years after the initial diagnosis. M12 cultures were established from a 75-year-old male patient who initially presented with a brain metastasis from a metastatic melanoma of unknown primary. MRI identified a paramedian mass in the right parietal region, while PET imaging revealed multiple extracranial metastases (highlighted by red circles). Similar to the findings with the M15 xenograft, both H&E-stained sections and Melan-A-stained sections of the M12 xenograft grown in mice displayed similar characteristics to the patient’s brain tumor. Notably, there was strong Melan-A/MLANA staining observed in both the patient's tumor sections and the corresponding M12 xenograft sections, demonstrating the consistency of the xenograft in maintaining one of the tumor's key features. The patient’s tumor responded well to treatment with vemurafenib, a BRAF inhibitor, followed by immunotherapy with ipilimumab and pembrolizumab. The patient has remained free of disease for more than ten years since the diagnosis. Scale bars, 100 µm. **(B)** Representative phase contrast microscopy images of normal human melanocytes (NHM) and M15 and M12 cultures are shown. NHM cells exhibit a spindled, elongated morphology characteristic of healthy melanocytes. In contrast, M12 cultures display a more varied morphology, with some elongated and some rounded cells, less distinct boundaries, and reduced dendritic processes. M15 melanoma cells are highly variable in shape and size, lacking the typical dendritic morphology and showing increased cell density and clustering, indicative of aggressive melanoma characteristics and a loss of normal melanocyte structure. Scale bars, 100 µm. **(C)** Quantitative PCR was used to assess gene expression from extracted mRNA of NHM, as well as M15 and M12 PDX cultures. The fold change in mRNA expression in M15 and M12 cultures relative to NHM is depicted. *MLANA*, Melan-A; *MITF*, Melanocyte Inducing Transcription Factor; *CDKN2A*, Cyclin Dependent Kinase Inhibitor 2A; *TP53*, Tumor Protein P53; *FN1*, Fibronectin; *TNC*, Tenascin C; *SPP1*, Secreted Phosphoprotein 1. **(D)** Cell surface integrin expression on NHM, as well as M15 and M12 PDX cultures, using flow cytometry. Fold change in integrin surface expression, normalized for isotype control background, compared to NHM is shown.

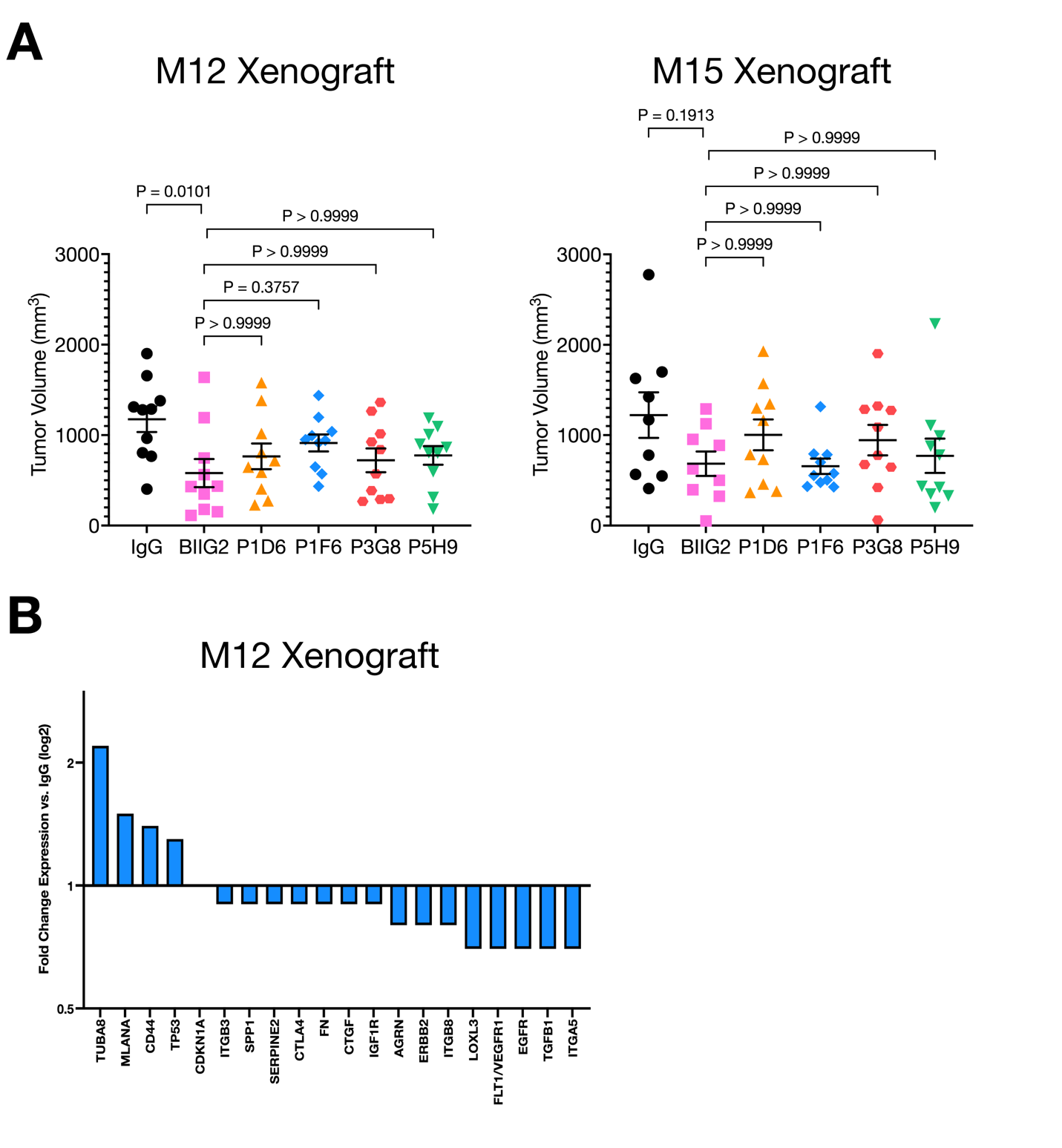

**Figure S5. Additional statistical analysis of BIIG2 efficacy and impacts of gene expression in M12 xenograft, Related to Figure 2. A)** Statistical analysis of BIIG2 effectiveness in reducing tumor volume against P1D6 and αv targeting antibodies. Data related to **Figure 2B**. Data are shown as individual tumor volume values with mean ± SEM; n=10 per antibody treatment, except for M15 where IgG control is n=9. A one-way ANOVA with a Bonferroni correction was performed to assess statistical significance of BIIG2 against the other antibodies. Adjusted P-values are shown and are considered significant if less than 0.05. **(B)** Analysis of tumors from M12 PDX showed that BIIG2 treatment of mice (n=9; 10 mg/kg twice weekly) induced the expression of the melanocyte differentiation marker Melan-A (MLANA) and cellular tumor antigen p53 (TP53) versus IgG (N=10) while inhibiting the expression of growth factors, growth factor receptors and the pro-angiogenic integrins (β3, β8 and α5).

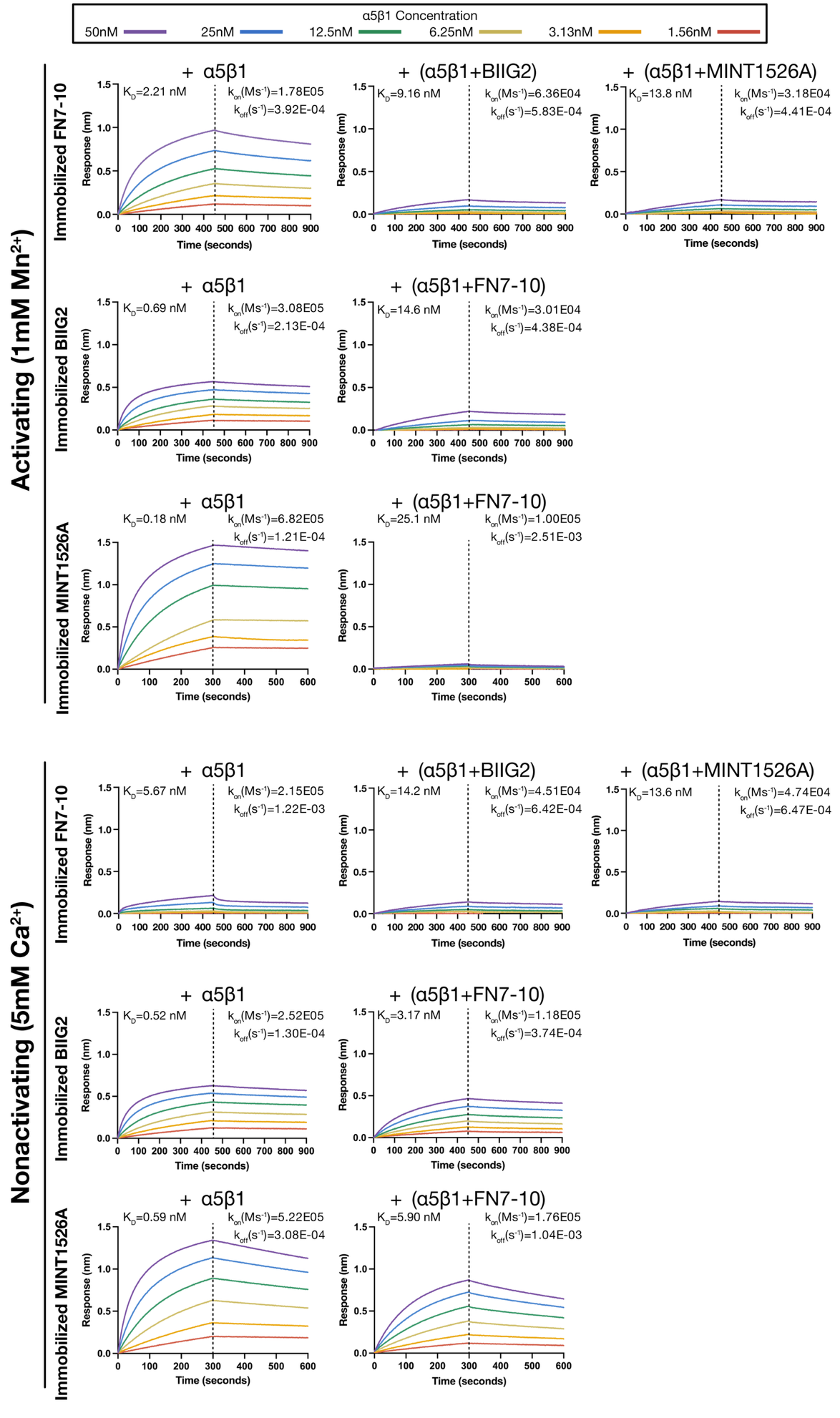

**Figure S6. Integrin and antibody BLI competition assay using a fragment of fibronectin (FN 7-10), Related to Figure 3.** Biolayer interferometry sensorgrams of competitive binding experiments. All experiments were loaded with 5 µg/mL of immobilized protein (BIIG2, MINT1526A, or FN 7-10). The color of the sensorgrams denotes the concentration of α5β1 ectodomain. The immobilized protein (BIIG2, MINT1526A, or FN 7-10) is denoted on the left and the titles of each sensorgram indicates the analyte (integrin α5β1 alone or pre-incubated integrin complex).

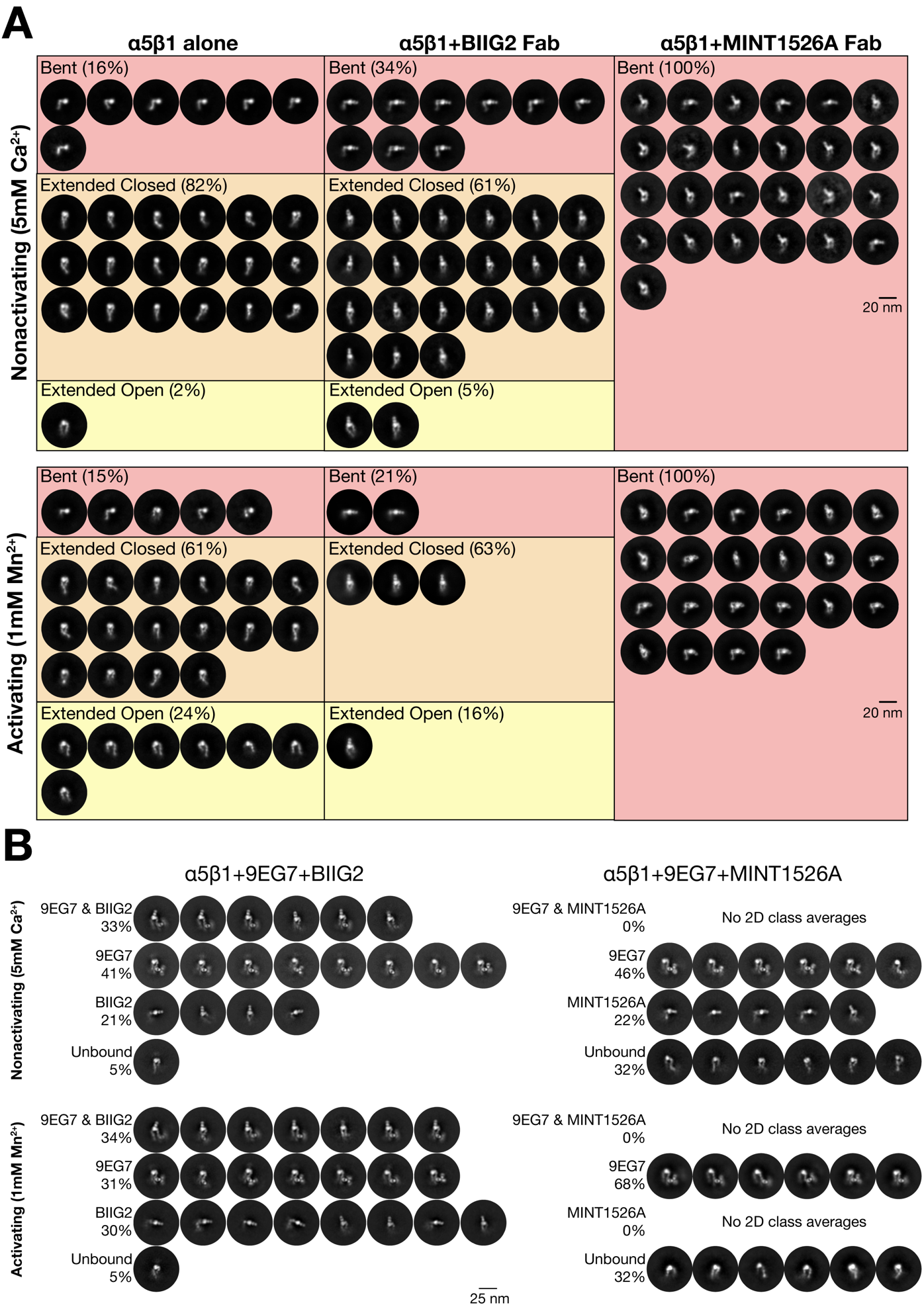

**Figure S7. Final nsEM 2D classes of apo α5β1, α5β1:BIIG2 Fab and α5β1:MINT1526A Fab, sorted by conformation and 2D classes of α5β1:9EG7:BIIG2 Fab and α5β1:9EG7:MINT1526A Fab complexes, Related to Figure 3.** **(A)** All negative stain EM 2D classes are shown for each condition evaluated to determine whether the binding of the Fabs influences α5β1 conformation. Scale bar is 20 nm. **(B)** Negative stain EM 2D classes of integrin α5β1 pre-incubated with 9EG7 antibody prior to addition of BIIG2 or MINT1526A Fabs. Scale bar is 25nm. Due to the varying particle numbers in each class and condition, the number of classes is not representative of the total number of particles contributing to that conformation; total particle percentages are listed.

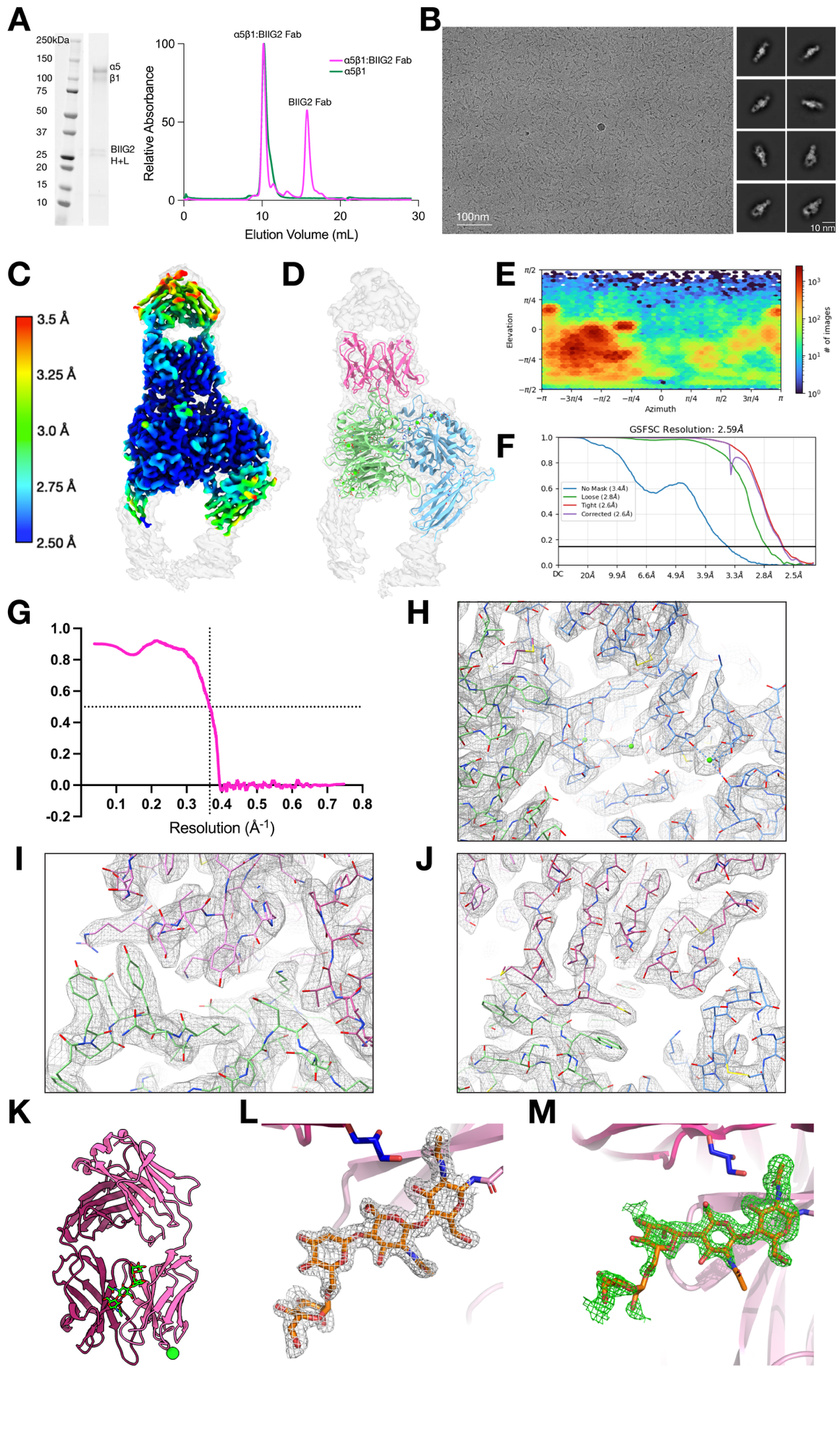

**Figure S8. BIIG2-α5β1 complex and structural supporting data, Related to Figure 4. (A)** SDS-PAGE of the first peak from Size Exclusion Chromatography (SEC) of the BIIG2:α5β1 complex, which contains both α5β1 and BIIG2 Fab (H, heavy chain; L, light chain fragments). SEC chromatogram of α5β1 alone (green) and α5β1:BIIG2 Fab complex (pink), with an excess of BIIG2 Fab. **(B)** Representative cryoEM micrograph and 2D class averages of the α5β1:BIIG2 Fab complex. Scale bars are 100 nm and 10 nm, respectively. **(C,D)** The locally refined final cryoEM map colored by resolution **(C)** or with the atomic model superimposed on the unsharpened map at a low threshold in semi-transparent white **(D)**. α5 is green; β1 is blue; BIIG2 is pink. **(E)** Angular distribution plot of particles used in the final cryoEM reconstruction. **(F)** Gold standard Fourier Shell Correlation (FSC). **(G)**: Map to model FSC. **(H-J)** Map and model showcasing clear density for key areas. The coordinated cations SyMBAS, MIDAS, and ADMIDAS **(H)**; the BIIG2 light-chain binding epitope, related to Figure 4D **(I)**, the BIIG2 heavy-chain binding epitope, related to Figure 4E **(J)** α5 is green; β1 is blue; BIIG2 is light-chain is lighter pink, BIIG2 is heavy-chain is darker pink, cations are neon green. **K)** Structure of antigen binding fragment of BIIG2 shown in cartoon representation (paratope on the bottom, light chain in light pink, heavy chain in dark pink) with ligands and solvent molecules depicted in stick representation. Four saccharide units of N-linked glycan N60 are shown in green stick representation and single chloride ion is shown as a green sphere. **(L)** Close-up view of N-linked glycan, with σ_A_-weighted 2*F*_o_-*F*_c_ electron density map (grey mesh, contoured at 1.0 σ) and N60 shown in stick representation. **(M)** Close-up view of N-linked glycan with polder (OMIT) map (green mesh, contoured at 3.0 σ) and N60 shown in stick representation. Protein structures and maps were generated using PyMOL (Schrödinger*,* Inc*.*) and UCSF ChimeraX.

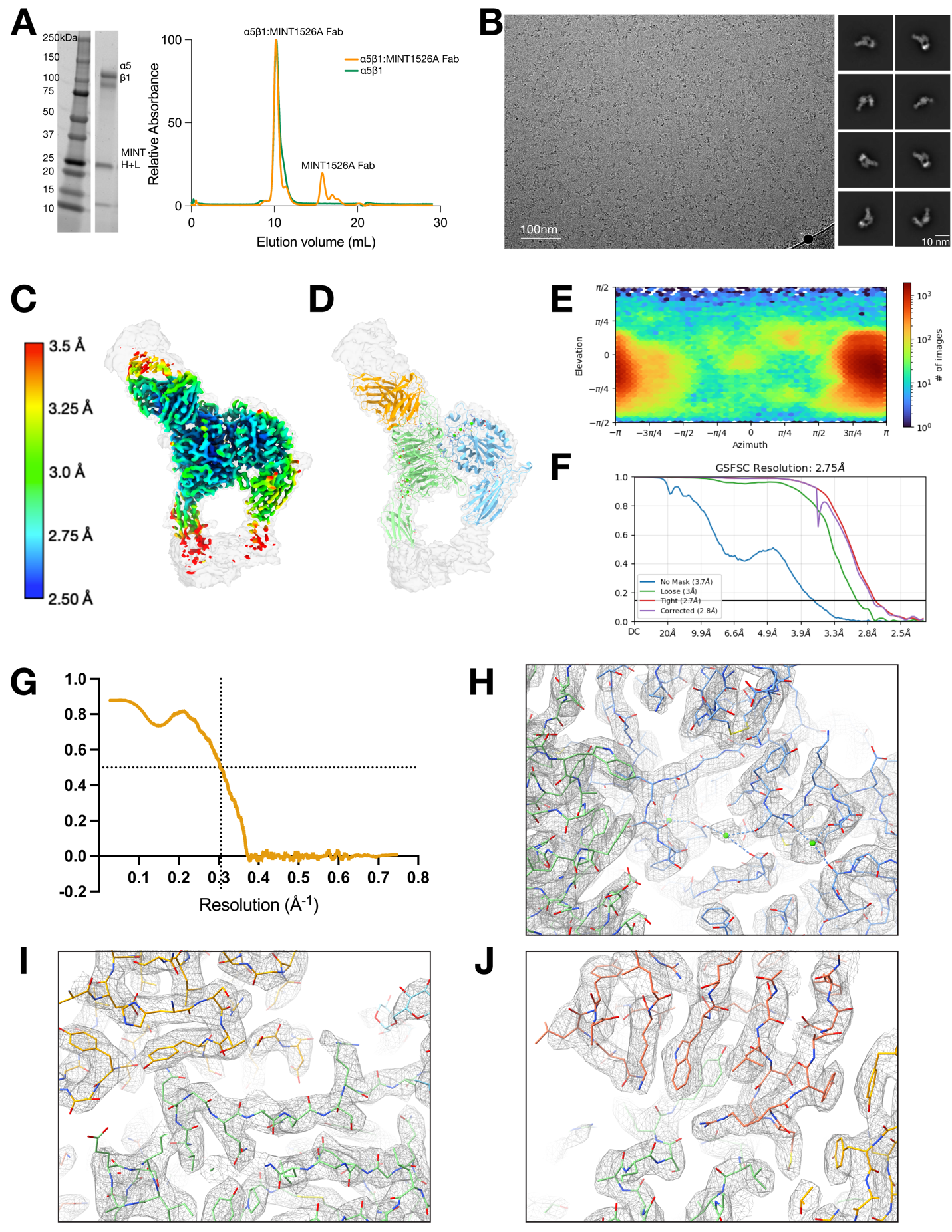

**Figure S9. MINT1526A-α5β1 complex and supporting cryoEM data, Related to Figure 5. (A)** SDS-PAGE of the first peak from Size Exclusion Chromatography (SEC) of the MINT1526A:α5β1 complex, which contains both α5β1 and MINT1526A Fab (H, heavy chain; L, light chain fragments). SEC chromatogram of α5β1 alone (green) and α5β1:MINT1526A Fab complex (orange), with an excess of MINT1526A Fab. **(B)** Representative cryoEM micrograph and 2D class averages of the α5β1:MINT1526A Fab complex. Scale bars are 100 nm and 10 nm, respectively. **(C,D)** The locally refined final cryoEM map colored by resolution **(C)** or with the atomic model superimposed on the unsharpened map at a low threshold in semi-transparent white **(D)**. α5 is green; β1 is blue; MINT1526A is orange. **(E)** Angular distribution plot of particles used in the final cryoEM reconstruction. **(F)** Gold standard Fourier Shell Correlation (FSC). **(G)**: Map to model FSC. **(H-J)** Map and model showcasing clear density for key areas. The coordinated cations SyMBAS, MIDAS, and ADMIDAS **(H)**; the MINT1526A heavy-chain binding epitope, related to Figure 5E. **(I)**, the MINT1526A light-chain binding epitope, related to Figure 5F **(J)** α5 is green; β1 is blue; MINT1526A is light-chain is lighter orange, BIIG2 is heavy-chain is darker orange, cations are neon green. Protein structure images and maps were generated using UCSF ChimeraX.

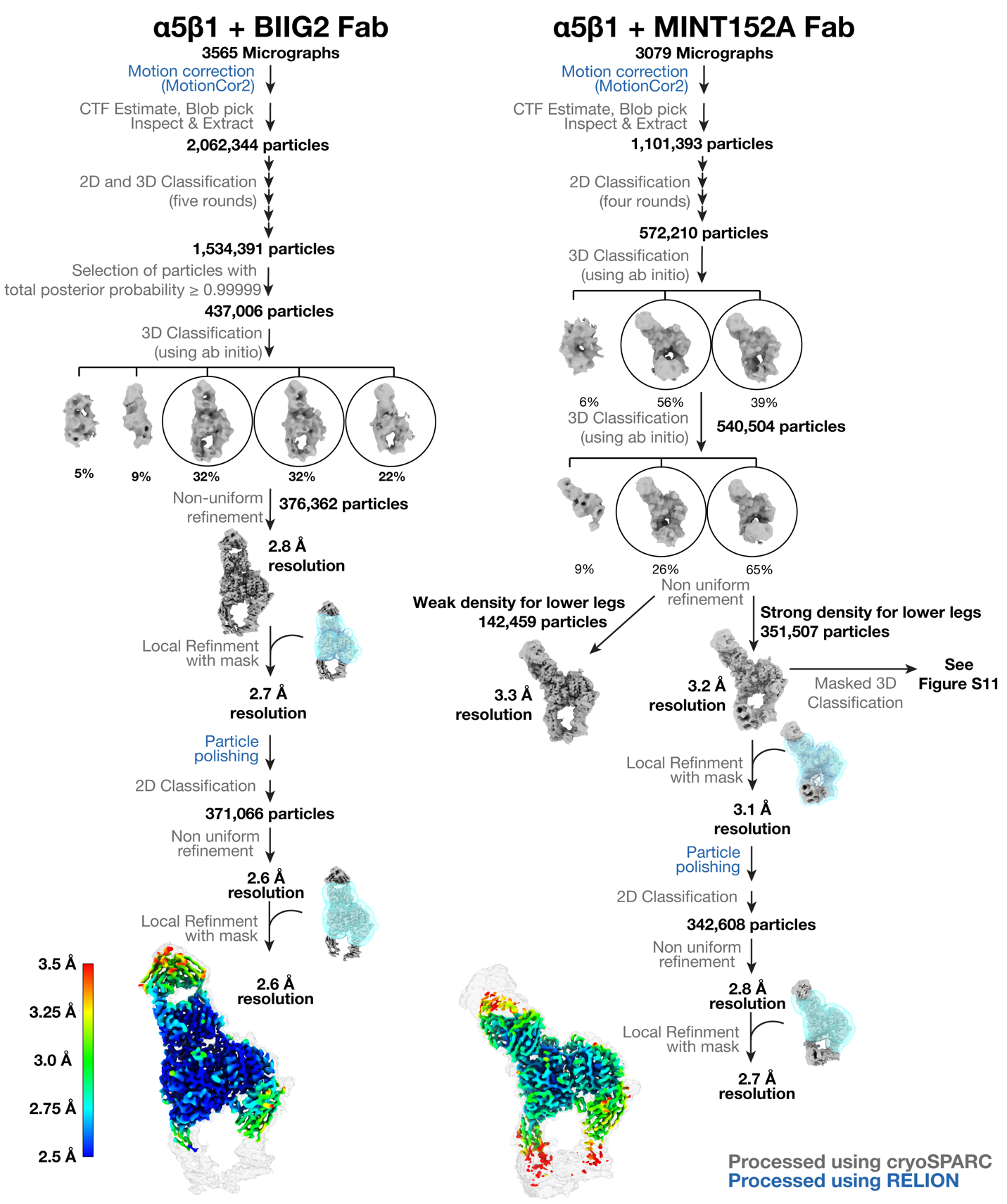

**Figure S10. CryoEM Processing Schematic of** **α5β1:BIIG2 Fab and α5β1:MINT1526A Fab, Related to STAR Methods.** A schematic flowchart highlighting the major classification and refinement steps for the α5β1:BIIG2 and α5β1:MINT1526A complexes. For each complex, a single dataset was collected. Particle numbers at significant steps are shown, and percentages of particles contributing to maps in 3D classification are shown. Intermediate maps are shown in gray and masks used for local refinement in semi-transparent blue. Final maps colored by local resolution at high threshold superimposed on a lower threshold map in semi-transparent white.

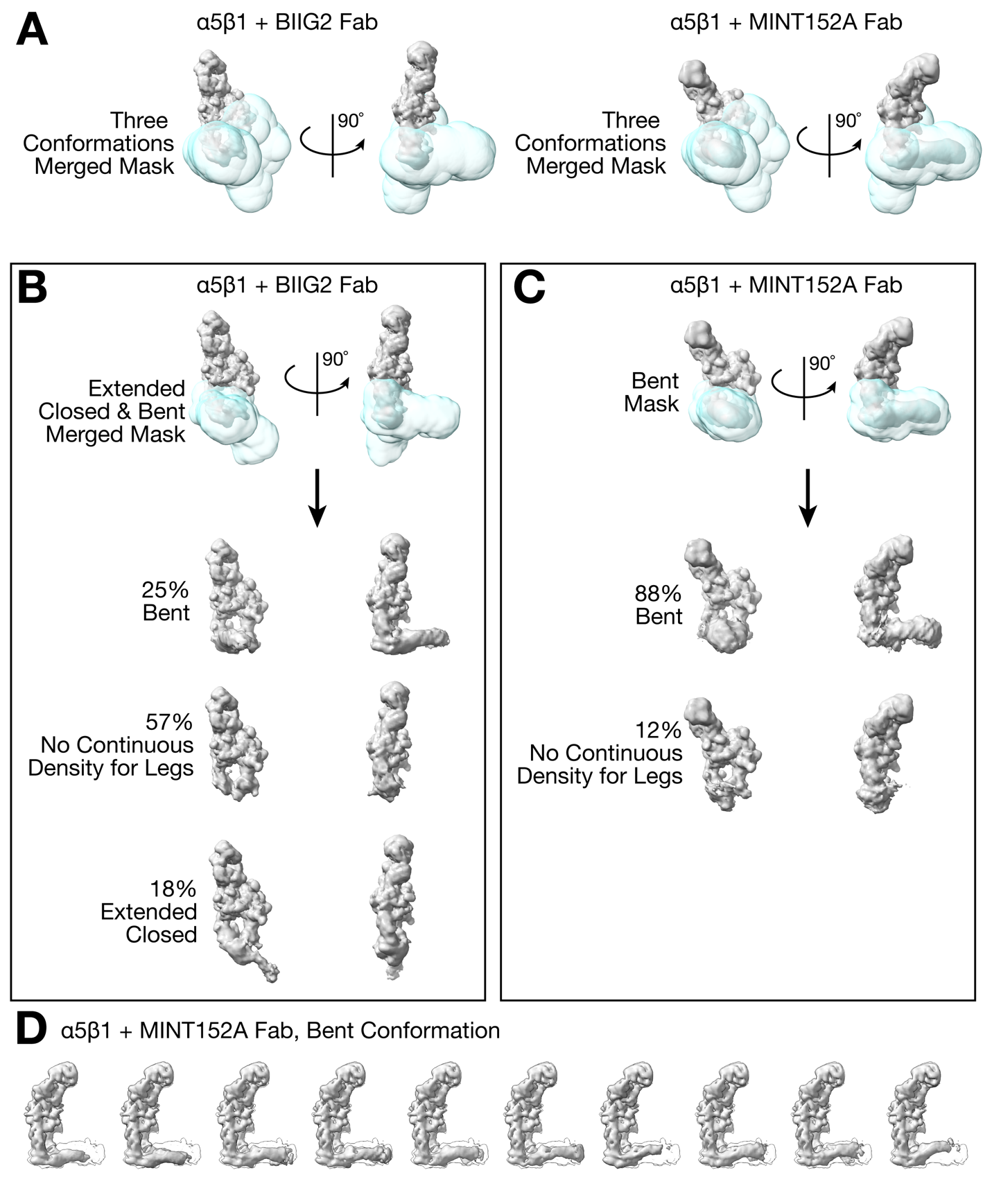

**Supplemental Figure S11. 3D classification defines the conformational states of integrin bound to BIIG2 or MINT1526A, Related to STAR Methods.** **(A)** First, the same soft, dilated mask encompassing the lower leg region was used for all three integrin conformations to determine which conformations were available when integrin was bound to either BIIG2 (left) or MINT1526A (right). The mask was made by merging the following maps: bent integrin (α5β1 + MINT1526A map, this work), extended open integrin (αvβ8, EMD-7939) ^57^, and extended open (α5β1, EMD-45655) ^83^. The BIIG2-bound dataset revealed lower legs in the bent and extended closed conformation, whereas the MINT1526A-bound dataset only showed lower legs in a bent conformation. **(B, C)** For a subsequent round of 3D classification, a mask encompassing the lower legs in a bent and extended closed conformation was used for BIIG2-bound integrin **(B)**; for MINT1526A-bound integrin, a mask encompassing the lower leg region in only the bent conformation was used **(C)**. Particles were sorted into twelve classes, and representative classes are shown. Masks are shown in semi-transparent cyan, maps in gray. **(D)** 3D classification illustrates the flexibility of the integrin lower legs of the α5β1:MINT1526A complex. The ten (of twelve) classes that showed strong density for the entire leg regions are shown. The integrin angle of rotation about the knee region ranges from 90 to 73 degrees, in agreement with previous data ^12^. Each panel depicts a single solid gray map overlaying the nine other maps shown in semi-transparent white.

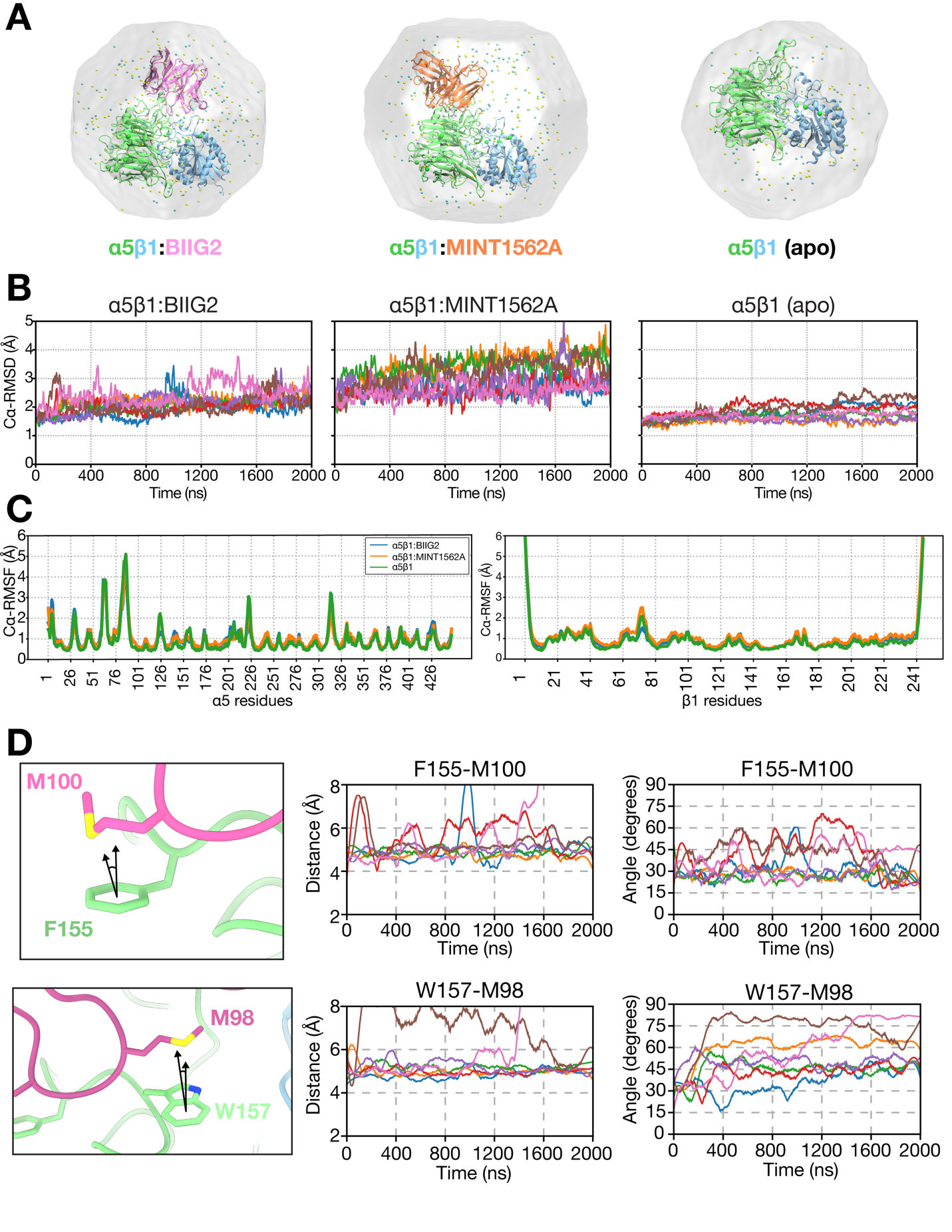

**Figure S12: Molecular dynamics simulations of apo α5β1 or in complex with BIIG2 and MINT1526A Fabs, Related to Figures 4 and 5. A)** Solvated α5β1:antibody MD system. The protein complexes were solvated in a truncated octahedron and NaCl ions. **(B)** Cα-RMSD of α5β1:antibody simulations. The seven MD replicates of 2000 ns are individually plotted. The initial structure was used as the reference structure for RMSD calculations. **(C)** Average per-residue Cα-RMSF of the α5 and β1 headpiece. Values represent average Cα-RMSF across seven 2000 ns replicates. **(D)** The location of the sulfur of the methionine-aromatic interactions between BIIG2 and α5β1. Distances are measured between the center of the aromatic ring of F155 (upper row) and W157 (lower row), and the center of the sulfur atoms. Arrows represent the normal vector of the aromatic plane and the angle measured. Distance and angle measurements in correlation with time, recorded with molecular dynamics (MD) simulations.

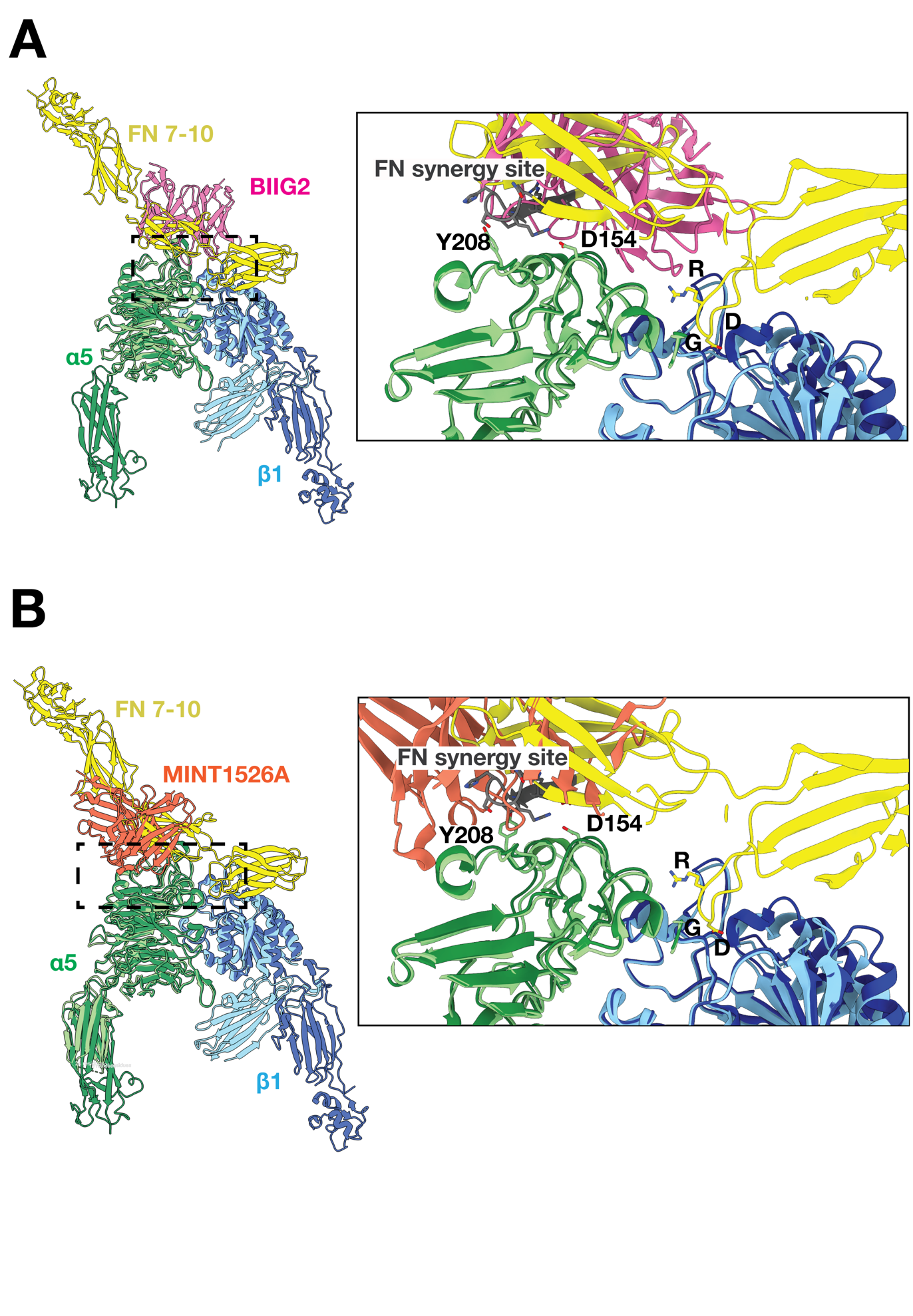

**Figure S13. BIIG2 and MINT1526A occlude Fibronectin binding, Related to Figure 4 and 5.** The previously determined structure (PDB ID: 7NWL) of integrin α5β1 (dark green, dark blue) bound to fibronectin domains 7-10 (yellow) is superimposed with either our structure of α5β1 (light green, light blue) bound to **(A)** BIIG2 fab (pink) or **(B)** MINT1526A fab (orange). The steric clash between fibronectin and BIIG2 occurs at the fibronectin type III domain 9; the clash between MINT1526A and fibronectin occurs at the fibronectin type III domain 9 and also blocks the synergy binding site.

**Table S1. Staining patterns of anti-human α5 integrin antibodies on normal FFPE tissue microarray (#BSB-0298), Related to STAR Methods.** PA5-82027, polyclonal rabbit α5 integrin antibody, antigen-verified; P1D6, monoclonal mouse α5 integrin antibody; BIIG2, monoclonal rat α5 integrin antibody.

| Tissue Type | PA5-82027 | P1D6 | BIIG2 |
| --- | --- | --- | --- |
| Chorionic Villi | +++ (Syncytioblast) | - | - |
| Breast | ++ (Ducts) | - | +++ (Fibroblasts) |
| Myometrium | +++ (Smooth Muscle) | - | +++ (Fibroblasts) |
| Cervix | - | - | +++ (Fibroblasts) |
| Fallopian Tube | - | + (Smooth Muscle) | +++ (Fibroblasts) |
| Brain | - | - | - |
| Stomach | + (Gastric Glands) | ++ (Gastric Glands) | +++ (Gastric Glands) |
| Adrenal Cortex | + (Cell Surface) | + (Cytoplasm) | ++ (Cytoplasm) |
| Pancreas | ++ (Acini) | - | ++ (Acini) |
| Salivary | ++ (Striated Ducts) | + (Striated Ducts) | + (Striated Ducts) |
| Colon | + (Smooth Muscle) | - | +++ (Fibroblasts) |
| Liver | +++ (Sinusoids) | +++ (Cords/Cytoplasm) | - |
| Kidney | + (Tubules) | + (Tubules) | + (Tubules) |
| Thyroid | - | - | - |
| Lung | ++ (Alveolar Ducts) | - | - |
| Skin | - | - | +++ (Fibroblasts) |
| Bladder | +++ (Smooth Muscle) | - | +++ (Fibroblasts) |
| Testis | - | - | - |
| Prostate | +++ (Fibromuscular Stroma) | - | - |
| Spleen | ++ (Germinal Center) | - | - |
| Tonsil | + (Germinal Center) | - | - |
| Bone Marrow | ++ (Megakaryocytes, Others) | - | +++ (Megakaryocytes, Others) |
| Thymus | - | - | - |

**Table S2. X-ray crystallography analysis of BIIG2 Fab fragment, Related to Figure 4.**

| Data Set | BIIG2 Fab^a^  (PDB 8R38) |
| --- | --- |
| **Data collection** |  |
| X-ray source | MAX-IV Laboratory - BioMAX |
| Wavelength, Å | 0.9537 |
| Space group | *P*2_1_2_1_2_1_ |
| Unit cell parameters – a,b,c (Å) | 62.9, 80.9, 87.5 |
| Resolution (Å)^b^ | 43.7 - 1.4 (1.43 - 1.40) |
| No. of unique reflections^b^ | 169,111 (8,404) |
| Multiplicity^b^ | 6.6 (4.5) |
| Completeness (%)^b^ | 99.6 (96.4) |
| Mean I/σ(I)^b^ | 7.59 (0.23) |
| CC_1/2_ (%)^b^ | 99.8 (10.4) |
| Wilson *B*-factor (Å^2^)^b^ | 21.3 |
| **Refinement** |  |
| *R*_work/_*R*_free_ (%)^c^ | 19.9 / 22.4 |
| Average *B* factors (Å^2^)  Protein  Ligand  Solvent | 26.1  38.0  32.9 |
| Number protein chains in the a.u. | 2 |
| Number of non-hydrogen atoms  Protein  Ligand  Solvent | 4,183  3,871  89  223 |
| r.m.s.d. from ideal geometry  Bond lengths (Å)  Bond angles (deg.) | 0.01  1.7 |
| Ramachandran plot  Favored (%)  Allowed (%)  Outliers (%) | 98.1  1.7  0.2 |

^a^Friedel pairs were treated as different reflections

^b^Values in parentheses refer to highest-resolution shell

^c^*R*_free_ was calculated from 5% of randomly selected reflections

**Table S3. Cryo-EM data collection, refinement, and validation statistics, Related to Figures 4 and 5.**

| Data Set | α5β1:BIIG2 Fab  (EMDB-44386)  (PDB 9B9J) | α5β1:MINT1526A Fab  (EMDB-44387)  (PDB 9B9K) |
| --- | --- | --- |
| **Data Collection and Processing** |  |  |
| Magnification | 36,000 | 36,000 |
| Voltage (kV) | 200 | 200 |
| Electron exposure (e^–^/Å^2^) | 50 | 50 |
| Defocus range (μm) | 1.4 – 1.8 (nominal) | 1.4 – 1.8 (nominal) |
| Pixel size (Å) | 1.122 | 1.122 |
| Symmetry imposed | C1 | C1 |
| Initial particle images (no.) | 2,425,530 | 1,101,393 |
| Final particle images (no.) | 371,066 | 342,608 |
| Map resolution (Å)           FSC threshold | 2.7  0.143 | 2.8  0.143 |
| Map resolution range (Å) | 2.5 – 6.5 | 2.6 - 8.0 |
| **Refinement** |  |  |
| Initial model used (PDB code) | 3VI3 (chains A,B)  BIIG2 Fab (this study, 8R38) | 7NXD (chains A,B)  MINT1526A Fab (this study) |
| Model resolution (Å)           FSC threshold | 2.7  0.5 | 3.3  0.5 |
| Model resolution range (Å) | 2.5 – 6.5 | 2.5 – 8.0 |
| Map sharpening *B* factor (Å^2^) | -50 | -75 |
| Model composition           Non-hydrogen atoms           Protein residues           Carbohydrates           Metal Ions | 8,518  1,055  32  7 | 9,150  1,142  32  7 |
| *B* factors (Å^2^)           Protein           Ligand | 93.3  N/A | 106.4  N/A |
| R.m.s deviations           Bond lengths (Å)           Bond angles (°) | 0.01  2.0 | 0.01  2.0 |
| Validation           MolProbity score           Clashscore           Poor rotamers (%) | 1.09  1.02  0.34 | 1.27  1.95  0.42 |
| Ramachandran plot           Favored (%)           Allowed (%)           Disallowed (%) | 95.4  4.2  0.4 | 95.7  4.3  0.0 |

**Table S4: Interactions of α5β1 with BIIG2 and MINT1526A Fabs, related to Figures 4 and 5.**

| α5β1:BIIG2 interactions | | | | | | |
| --- | --- | --- | --- | --- | --- | --- |
| α5β1 Integrin | | | BIIG2 Fab | | | Interaction Type |
| Residue | | Subunit | Residue | | CDR |  |
| E | 81 | alpha | K | 27 (L) | CDRL1 | Salt bridge |
| E | 124 | alpha | K | 92 (L) | CRDL3 | Salt bridge |
| E | 124 | alpha | Y | 94 (L) | CDRL3 | Hydrogen bond |
| K | 125 | alpha | H | 91 (L) | CDRL3 | Hydrogen bond |
| D | 154 | alpha | K | 31 (L) | CDRL1 | Hydrogen bond |
| D | 154 | alpha | Y | 32 (L) | CDRL1 | Hydrogen bond |
| Y | 205 | alpha | R | 28 (L) | CDRL1 | Hydrogen bond |
| Y | 208 | alpha | S | 30 (L) | CDRL1 | Hydrogen bond |
| E | 125 | alpha | D | 95 (H) | CDRH3 | Hydrogen bond |
| E | 125 | alpha | R | 96 (H) | CDRH3 | Hydrogen bond |
| E | 126 | alpha | T | 97 (H) | CDRH3 | Hydrogen bond |
| E | 126 | alpha | N | 33 (H) | CDRH1 | Hydrogen bond |
| D | 154 | alpha | T | 100A (H) | CDRH3 | Hydrogen bond |
| S | 156 | alpha | G | 99 (H) | CDRH3 | Hydrogen bond |
| S | 189 | Beta | G | 54 (H) | CDRH2 | Hydrogen bond |
| E | 190 | Beta | R | 71 (H) | Framework | Salt bridge |
| α5β1:MINT1526A interactions | | | | | | |
| α5β1 Integrin | | | MINT1526A Fab | | | Interaction Type |
| Residue | | Subunit | Residue | | CDR |  |
| R | 144 | alpha | P | 52C (H) | CDRH2 | Hydrogen bond |
| E | 202 | alpha | K | 52 (H) | CDRH2 | Hydrogen bond |
| E | 202 | alpha | R | 99 (H) | CDRH3 | Hydrogen bond |
| S | 203 | alpha | R | 99 (H) | CDRH3 | Hydrogen bond |
| L | 212 | alpha | R | 99 (H) | CDRH3 | Hydrogen bond |
| Y | 204 | alpha | N | 53 (H) | CDRH2 | Hydrogen bond |
| Y | 205 | alpha | W | 33 (H) | CDRH1 | Hydrogen bond |
| Y | 205 | alpha | L | 95 (H) | CDRH3 | Hydrogen bond |
| E | 207 | alpha | R | 32 (H) | CDRH1 | Hydrogen bond |
| Y | 208 | alpha | R | 32 (H) | CDRH1 | Hydrogen bond |
| Y | 208 | alpha | Y | 102 (H) | CDRH3 | Hydrogen bond |
| L | 212 | alpha | G | 97 (H) | CDRH3 | Hydrogen bond |
| Q | 214 | alpha | T | 32 (L) | CDRL1/ Framework | Hydrogen bond |
| Q | 214 | alpha | S | 91 (L) | CDRL3 | Hydrogen bond |
| R | 220 | alpha | D | 52A (L) | CDRL2 | Hydrogen bond |

**Table S5. Primer list for cDNA amplification by quantitative PCR and α5β1 ectodomain construct design, Related to STAR Methods.**

| **Primer Name** | **Sequence** |
| --- | --- |
| ACTB(1)-forward | GGCTACAGCTTCACCACCAC |
| ACTB(1)-reverse | TAATGTCACGCACGATTTCC |
| ACTB(2)-forward | TGCTATCCCTGTACGCCTCT |
| ACTB(2)-reverse | GAGTCCATCACGATGCCAGT |
| ACTB(3)-forward | GGACTTCGAGCAAGAGATGG |
| ACTB(3)-reverse | CTTCTCCAGGGAGGAGCTG |
| RPL8(1)-forward | ACAGAGCTGTGGTTGGTGTG |
| RPL8(1)-reverse: | TTGTCAATTCGGCCACCT |
| RPL8(2)-forward | ACTGCTGGCCACGAGTACG |
| RPL8(2)-reverse | ATGCTCCACAGGATTCATGG |
| RPLP0(1)-forward | AACTCTGCATTCTCGCTTCC |
| RPLP0(1)-reverse | GCAGACAGACACTGGCAACA |
| RPLP0(2)-forward | GCACCATTGAAATCCTGAGTG |
| RPLP0(2)-reverse | GCTCCCACTTTGTCTCCAGT |
| MITF-forward | CGGCATTTGTTGCTCAGAAT |
| MITF-reverse | GAGCCTGCATTTCAAGTTCC |
| MLANA-forward | AGAGAAAAACTGTGAACCTGTGG |
| MLANA-reverse | ATAAGCAGGTGGAGCATTGG |
| FN1-forward | GAAGTGGTCATTTCAGATGTGATT |
| FN1-reverse | CCATTGTCATGGCACCATCT |
| SPP1(1)-forward | TGTGGGGCTAGGAGATTCTG |
| SPP1(1)-reverse | GGCTAAACCCTGACCCATCT |
| SPP1(2)-forward | GTTTCGCAGACCTGACATCC |
| SPP1(2)-reverse | TCCTCGTCTGTAGCATCAGG |
| TNC-forward | GGTACAGTGGGACAGCAGGT |
| TNC-reverse | GATCTGCCATTGTGGTAGGC |
| CDKN2A-forward | AACGCACCGAATAGTTACGG |
| CDKN2A-reverse | CATCATCATGACCTGGATCG |
| OpenVect_for_a5_fwd | AGGCAGCTATGGAACAGGAGGCCTGGAGGT |
| OpenVect_for_a5_rev | GGCTTCCCATGGTGGCAAGCTTAAGTTTAAACGCTAG |
| alhpa5insert_fwd | GCTTGCCACCATGGGAAGCCGGACGCCAGA |
| alpha5insert_rev | CTCCTGTTCCATAGCTGCCTTCTGCCTTGGTCC |
| Correct_a5_fwd | GGAACAGGAGGCCTGGAGGTG |
| Correct_a5_rev | ATAGCTGCCTTCTGCCTTG |
| OpenVect_for_b1_fwd | TGGTCCAGACGATACCTCTGGCCTGGAGGTG |
| OpenVect_for_b1_rev | GTAAATTCATGGTGGCAAGCTTAAGTTTAAACGC |
| beta1insert_fwd | GCTTGCCACCATGAATTTACAACCAATCTTCTGGATTGG |
| beta1insert_rev | CAGAGGTATCGTCTGGACCAGTGGGACACTC |
